# Integrating gene mutation spectra from tumors and the general population with gene expression topological networks to identify novel cancer driver genes

**DOI:** 10.1101/2023.05.02.539093

**Authors:** Dan He, Ling Li, Zhiya Lu, Shaoying Li, Tianjun Lan, Feiyi Liu, Huasong Zhang, Bingxi Lei, David N. Cooper, Huiying Zhao

## Abstract

**Background:** Understanding the genetics underlying cancer development and progression is the most important goal of biomedical research to improve patient survival rates. Recently, researchers have proposed computationally combining the mutational burden with biological networks as a novel means to identify cancer driver genes. However, these approaches treated all mutations as having the same functional impact on genes and incorporated gene-gene interaction networks without considering tissue specificity, which may have hampered our ability to identify novel cancer drivers.

**Methods:** We have developed a framework, DGAT-cancer that integrates the predicted pathogenicity of somatic mutation in cancers and germline variants in the healthy population, with topological networks of gene expression in tumor tissues, and the gene expression levels in tumor and paracancerous tissues in predicting cancer drivers. These features were filtered by an unsupervised approach, Laplacian selection, and those selected were combined by Hotelling and Box-Cox transformations to score genes. Finally, the scored genes were subjected to Gibbs sampling to determine the probability that a given gene is a cancer driver.

**Results:** This method was applied to nine types of cancer, and achieved the best area under the precision-recall curve compared to three commonly used methods, leading to the identification of 571 novel cancer drivers. One of the top genes, *EEF1A1* was experimentally confirmed as a cancer driver of glioma. Knockdown of *EEF1A1* led to a ~ 41-50% decrease in glioma size and improved the temozolomide sensitivity of glioma cells.

**Conclusion:** By combining the pathogenic status of mutational spectra in tumors alongside the spectrum of variation in the healthy population, with gene expression in both tumors and paracancerous tissues, DGAT-cancer has significantly improved our ability to detect novel cancer driver genes.

## Background

The identification of cancer driver genes is important for the early diagnosis of cancer, for identifying efficacious anti-cancer therapeutics and for investigating the underlying mechanisms of tumorigenesis. Traditionally, cancer driver genes have been recognized on the basis of their being recurrently altered in tumors[1, 2]. However, the ability of many somatic mutations to alter gene function is often uncertain, making it hard to identify cancer driver genes unambiguously. Clearly, we require information other than somatic mutation data in order to reliably detect cancer driver genes.

Traditional approaches to identifying cancer driver genes have relied upon the statistical testing of the mutational burden of individual genes[1, 3], a strategy that assumes that driver mutations occur more frequently than expected by chance alone[4]. This type of approach, exemplified by MutSigCV[4] and MuSic[5], is in common use and has been successful in identifying many genes that harbor recurrent mutations in cancer(s). Such approaches are useful for identifying those genes which are mutated across a large number of samples but can easily miss genes mutated in only a small number of samples. However, most cancer driver genes are only mutated in a small proportion of patients[6]. Indeed, somatic mutation frequencies are influenced by specific characteristics of the gene, such as length, replication time and mutation rate in the healthy population[7–10]. A high gene mutation rate in the healthy population suggests that many somatic mutations within that gene are likely to be neutral passengers rather than drivers[11, 12]. Various other methods have been developed that improve our ability to recognize genes characterized by a higher-than-usual rate of somatic mutation[4, 5, 11–13]. OncodriveFML[14] is one such method designed to use the patterns of mutations across tumors in coding and non-coding regions to identify cancer driver genes. Similarly, OncodriveCLUSTL[15] is a method that detects cancer driver genes by clustering the mutations in cancer cohorts according to the number of the mutation distribution.

Mutations residing within transcribed regions of the genome are known to be more likely to influence the gene expression profiles of tumors[4]. The mutation rates of specific genes can be cross-compared with tumor expression signatures[16, 17]. Cancer driver genes could in principle be identified by integrating gene mutation data with gene expression data. As an example, a previous study performed enrichment analysis to integrate genomic and transcriptomic alterations from whole-exomes and functional data from protein function predictions with gene interaction networks to reveal breast cancer driver genes[18]. Recent approaches have been developed by combining mutation scores with biological network protein-protein interaction (PPI) data to predict cancer driver genes[19, 20]. These approaches did not consider functional data for mutations and cancer tissue-specific PPI networks, which may reduce the ability in predicting novel cancer drivers. Thus, no approach to identifying cancer driver genes has yet been devised that fully considers the pathogenic status of mutational spectra in tumors alongside the spectrum of variation in the healthy population, in combination with gene expression in both tumors and normal tissues.

To address these shortcomings, we devised a new model, DGAT-cancer (Distinguish cancer drivers using Genomics and Transcriptome data), which integrates the predicted pathogenicity scores of somatic mutations in cancers and germline mutations in the healthy population, with gene expression in tumors and paracancerous tissues in order to detect cancer driver genes. The work scheme of DGAT-cancer is shown in Fig. 1. First, for each gene, DGAT-cancer calculated a unidirectional Earth Mover’s Difference score (uEMD) to evaluate the difference between the predicted pathogenic scores (obtained from 19 predictors in dbNSFP[21]) of somatic mutational spectra in cancers and that of germline variants in healthy populations. The influence of mutation on gene expression has been evaluated by integrating gene expression data in tumors with somatic mutations through topological data analysis (TDA)[22]. Briefly, using TDA, we have constructed a gene expression topological network by clustering samples with similar gene expression profiles. The gene expression topological network was then used to evaluate the frequencies of mutational spectra occurring in different sample clusters. For those mutations occurring in adjacent sample clusters, we calculated the divergence of their frequencies across clusters, which was used as one feature of the gene in which the mutations resided. The Jensen–Shannon divergence between the gene expression profile and mutation profile of each gene on the topological network was further computed and used as another feature of the gene. In total, 24 features for each gene were included in DGAT-cancer. The most effective features were identified by Laplacian selection in an unsupervised way. The selected features were integrated by means of the Hotelling and Box-Cox transformations to score the genes. Finally, by using gene scores as weights, we performed Gibbs sampling to identify cancer drivers. This method was then applied to 6,643 samples containing mutation and gene expression data from tumors and paracancerous tissues derived from 9 cancer cohorts and succeeded in identifying 734 genes as being significant cancer drivers. Of these, 571 were previously unreported as cancer-associated genes. Further, these genes were found to be highly enriched in pathways related to cancer, as well as significantly enriched in drug-response genes, demonstrating that this new approach facilitates the identification of clinically relevant genes. One of the top genes, *EEF1A1,* was predicted to be a driver of glioma. This is the first time that *EEF1A1* has been considered to be a driver of glioma. Its relationship with glioma was confirmed by analysis of surgical specimens, a cell model and an animal model.

**Fig. 1.**
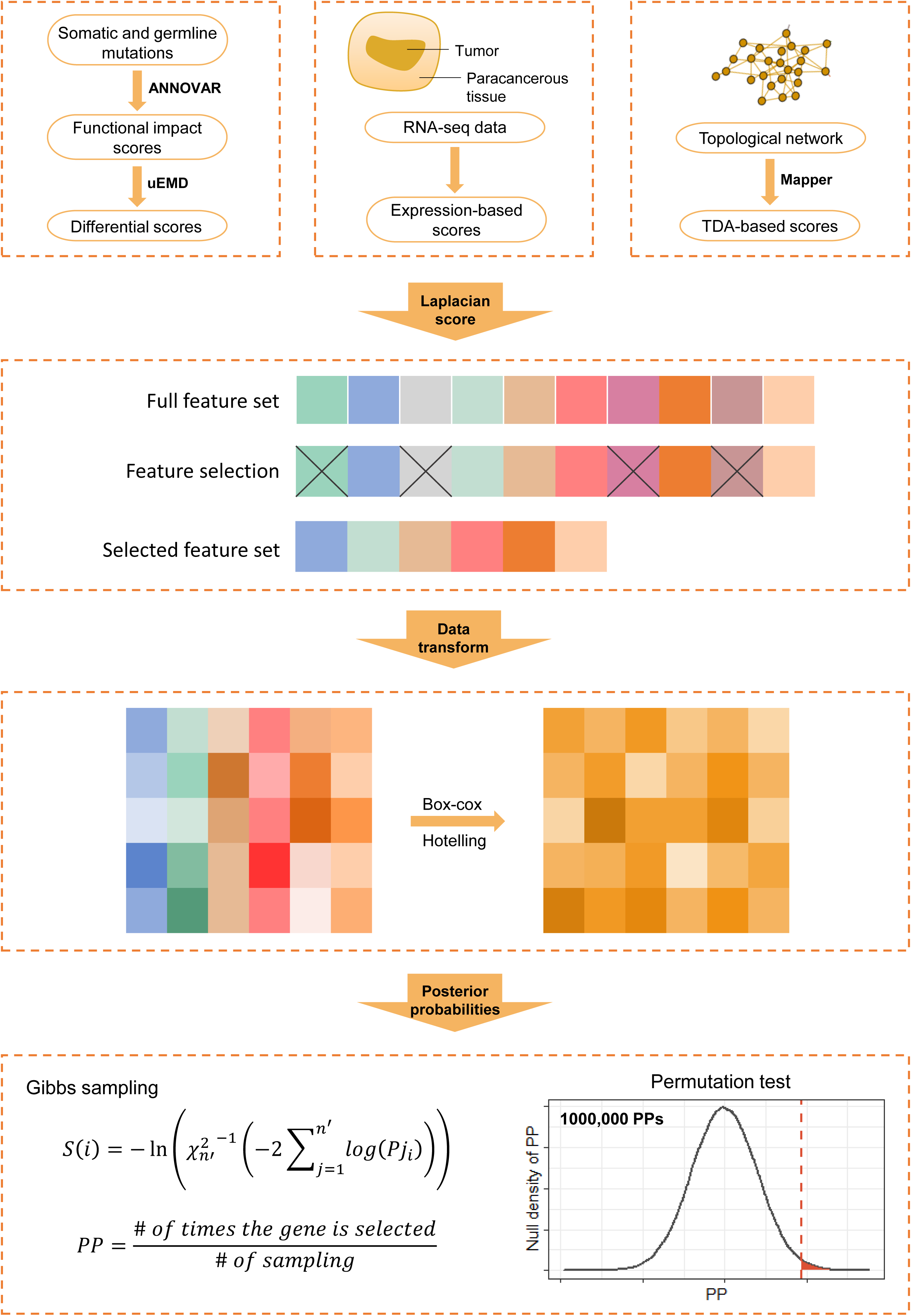
Overview of DGAT-cancer. First, somatic mutations were collected from two databases, ICGC and TCGA, as well as germline mutations from the 1000 Genomes Project. The predicted pathogenicity scores of these mutations were generated by ANNOVAR. The differences between the pathogenicity scores of mutated genes in cancer (somatic) and those in the general population (germline) were evaluated by DGAT-cancer calculating the uEMD scores. RNA-seq data were collected from tumors and paracancerous tissues. The RNA-seq data from tumors were used to construct a topological network. The gene expression topological network was then used to evaluate the frequencies of mutation spectra occurring in different sample clusters. For the mutations occurring in adjacent sample clusters, we calculated the divergence of their frequencies across clusters, which was used as one feature of the genes harboring the mutations. We also calculated the Jensen–Shannon divergence between the mutation frequency and the mRNA expression of the gene across clusters of the network as one feature. All collected features of genes were filtered using Laplacian scores. We transformed these selected features using Hotelling and Box-Cox transformations in order to integrate them into one composite risk score. By using gene scores as weights, we performed Gibbs sampling to sample genes in order to identify cancer drivers. Finally, we generated null distributions of PP for each gene to compute an experience P-value. The genes with *p_adj_* < 0.01 were selected as candidate cancer driver genes.

## Methods

### Data collection

#### Somatic mutation data

We collected simple somatic mutation (SSM) data from 12 cancer cohorts from the Broad Institute GDAC Firehose Portal, the International Cancer Genome Consortium (ICGC) Data Portal and The Cancer Genome Atlas (TCGA) (Additional file 1: Table S1). The coordinates of the data are by reference to the genome assembly version hg19. The germline mutation data of 2,557 individual samples were collected from Phase 3 of the 1000 Genomes Project (GRCh38). Although gnomAD[23] contains germline variants from a larger cohort, it does not provide genetic data from each sample and was therefore inappropriate for use in generating mutation score profiles for this study. Somatic mutation entries that were duplicated between multiple databases were removed such that only one non-redundant entry was retained.

#### The pathogenicity of mutations

There are many methods available with which to predict the pathogenicity of mutations. We employed dbNSFP (version: dbnsfp30a, also called LJB*)[21] of ANNOVAR[24] to annotate mutations from the 1000 Genomes Project (1000GP) and the somatic mutations (Additional file 1: Table S1). The dbNSFP includes 19 predictors which provide scores representing the probability of the non-synonymous variants being pathogenic (Additional file 2: Table S2). The scores given by each predictor were normalized into the range of 0 to1 in this study.

#### Known cancer driver genes

The accuracy of the predictions made by DGAT-cancer was examined using a set of known cancer driver genes. This set of 1,168 unique cancer driver genes comprised a non-redundant set of 723 genes from the COSMIC Cancer Gene Census[25] (CGC) and 1,064 genes from OncoKB[26]. OncoKB contains more cancer driver genes than CGC because it collects cancer genes from various panels, including class Tier 1 genes in CGC (576), the MSK-IMPACT™ panel (505), the MSK-IMPACT™ Heme and HemePACT panels (575), the FoundationOne CDx panel (324), the FoundationOne Heme panel (593), and data from Vogelstein et al.(125)[6]. We also collated gene sets containing cancer type-specific driver genes, which were collected from IntOGen[27]. The number of known driver genes for each cancer type are shown in Additional file 1: Table S3. These genes were used as gold standard cancer-associated genes to evaluate the accuracy of the predictions made in this study.

#### Constrained genes

A previous study[28] provided a set of 1,003 genes that are significantly lacking in missense variations (NHLBI’s Exome Sequencing Project), suggesting that they have a high intolerance to germline mutation.

#### Genes affecting cell proliferation and/or viability

These genes were identified by a previous study that performed shRNA screens in 216 cancer cell lines[29]. We downloaded the final shRNA quality file from the study and selected 687 genes as significantly affecting cell survival if the genes had shown significant gene suppression in cells according to the evaluation of ATARiS[30] (*q* < 0.05) in at least one half of the shRNA screens.

#### Drug response genes

From OncoKB[26], we collected 43 genes that have been reported to respond to FDA-approved drugs (Level 1) and 17 genes that respond to drugs used in standard care (Level 2). From IntOGen[27], we identified 51 genes whose protein products interact with FDA-approved drugs[31]. From these datasets, we removed off-target genes, gene therapy targets in the IntOGen list, and drugs targeting fusion driver genes, as well as 15 genes associated with drug resistance in COSMIC[32]. Finally, 46 actionable genes were obtained which could be used for drug target enrichment analysis.

#### Pathology data

Genes whose mRNA expression significantly correlated with the survival of cancer patients were downloaded from The Human Protein Atlas (HPA)[33]. The HPA performed Kaplan-Meier analysis to estimate the correlation between mRNA expression and patient survival for each gene (a total of 20,090 genes) for 20 cancer types. We selected genes with expression levels significantly (log-rank *p* < 0.05) correlating with patient survival in at least one cancer type investigated by this study. The proportion of genes predicted to be cancer drivers by DGAT-cancer in these cancer survival-related genes was compared to those genes predicted to be non-cancer drivers by a one-sided Fisher’s Exact test.

### Pathogenicity score profiles of mutations in cancers and in the healthy population cohort

Each tumor sample or specific individual from the 1000GP may harbor more than one mutation in each gene. The pathogenic status of these mutations was scored by 19 distinct approaches each representing different detrimental effects of mutations on gene function. The mutation score of a given gene for a specific sample determined by a particular predictive approach was defined as the average mutation score across all mutations in that gene. Mutation scores of a given gene from all tumor samples or all samples from 1000GP (Additional file 1: Table S1) were considered to represent the mutation score profiles of that gene calculated by one particular predictive approach. Then, for each gene, we constructed a density distribution for mutation scores in each cancer cohort and in the healthy population cohort, respectively. The density distribution was divided into 100 evenly spaced bins. A gene was filtered out from the construction of mutation score profiles if all the mutations in the gene were derived from fewer than five samples.

In order to compare the mutation score profiles in tumors with that of the healthy population cohort, we calculated the difference between the two profiles by means of a unidirectional Earth Mover’s Difference score (uEMD, *Equation (1)*). For each gene, we repeated this calculation for all 19 types of functional score and obtained 19 uEMD scores. These uEMD scores were termed uEMD-Mut scores.

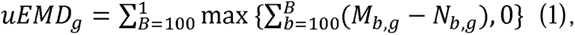

where *M_b,g_* is the fraction of normalized scores in the *b*-th bin for gene *g* in the density distribution of a cancer cohort, *N_b,g_* is the fraction of normalized scores in the *b*-th bin for gene *g* in the density distribution of the general population, and *B* is the index of bins in the density distribution. Thus, genes with a significantly different distribution in cancer tissues from the general cohort would be given higher uEMD scores.

### Comparing gene expression profiles of tumors and paracancerous tissues

RNA-seq data (RSEM normalized count, log2 transformed) of tumors from 12 cancer types in TCGA were downloaded from the UCSC Xena platform[34]. The corresponding RNA-seq data of paracancerous tissues from eight types of cancer including Bladder urothelial carcinoma (BLCA), Breast invasive carcinoma (BRCA), Cervical and endocervical cancers (CESC), Colon adenocarcinoma (COAD), Glioblastoma multiforme (GBM), Head and neck squamous cell carcinoma (HNSC), Lung adenocarcinoma (LUAD) and Stomach adenocarcinoma (STAD) were downloaded from TCGA. RNA-seq data from paracancerous tissues or tumors with a sample size fewer than five were not included in the study. The sample sizes of RNA-seq data for each type of cancer are shown in Additional file 1: Table S4. For each gene, we calculated the median expression level across samples. Then, we computed the uEMD score for each gene to measure the difference in the density distribution of gene expression in tumor and paracancerous tissues. This uEMD score of the gene was termed uEMD-Ex.

### Topological network constructed from gene expression data and somatic mutations in tumors

By using gene expression data derived from tumor samples, we constructed a topological network for each type of cancer (Additional file 1: Table S4 and Table S5) using Mapper algorithm, an R package in TDAmapper (the detailed input arguments are listed in Additional file 1: Table S6)[35]. Briefly, tumor samples with similar expression profiles are clustered into one node. Where two nodes have at least one tumor sample in common, they are connected by an edge. For each gene in the topological network, we evaluated the divergence of the mutation frequency of the gene in samples across similar gene-expression clusters (*Equation (2)*). The strategy used here is similar to that described in previous studies[22, 36]. Prior to the calculation, we used a threshold of MAF > 0.001 to filter out mutations occurring at low frequencies in tumors.

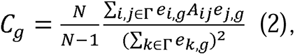

where Γ denotes the set of nodes in the topological network, *A* is the adjacency matrix of the topological network, *N* is the number of nodes in Γ, and *e_i,g_* is the average frequency of non-synonymous mutations of gene *g* for samples in the node *i*. A lower *C_g_* represents a higher divergence of the mutation frequency between similar clusters.

To evaluate the similarity between the profiles of mutation frequency and mRNA expression, we computed the Jensen–Shannon divergence between the expression and mutation profiles of each gene based on the topological network (*Equation (3)*). *C_g_* and *JSD_g_* were termed MutExTDA scores.

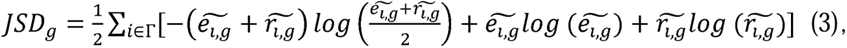

where 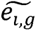 denotes the fraction of tumors with gene *g* somatically mutated in node *i*, 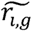 denotes the average expression of gene *g* in the tumors associated with node *i*, and Γ denotes the set of nodes in the topological network. Prior to calculation, they were normalized to meet 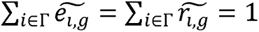. A lower *JSD_g_* denotes less difference between the two distributions.

### Data preprocessing

We obtained 24 features to describe each gene, comprising 19 uEMD-Mut scores, one uEMD-Ex score, two MutExTDA scores and two features representing the median expression levels of the gene in tumor and paracancerous tissues, respectively (Additional file 2: Table S2). We filtered out genes lacking more than half the number of features (Additional file 3: Table S7 and Additional file 1: Table S8). For the remaining genes, we used k-nearest neighbors (KNN) imputation to fill in the missing features. More than 99% of genes for cancer types Pheochromocytoma and paraganglioma (PCPG), Testicular germ cell tumors (TGCT) and Thyroid carcinoma (THCA) were lacking uEMD-Mut scores. For these cancer types, we constructed the predictor only using gene expression-based features (uEMD-Ex or MutExTDA) to perform the prediction.

### Using Laplacian Score to select features

We used an unsupervised method, the Laplacian Score, to select features that have high power to preserve the local geometric structure of the feature space[37]. The detailed steps for the application of the Laplacian Score in feature selection were as follows:

1. A network *N* was constructed to connect all *m* candidate genes (*N* ∈ ℝ*^m×m^*). For each pair of genes, we calculated the Euclidean distance, |*x_i_* − *x_j_|* between their feature vectors (*x_i_* is the feature vector of gene *i*, and *x_j_* is the feature vector of gene *j*). Based on the Euclidean distance, if gene *i* is among the top *k* (here we set *k* = [0.01 × *m*]) of the nearest genes to gene *j,* or the gene *j* is among the top *k* of the nearest gene to the gene *i* (*i ≠ j*), *we* set the connection between *i* and *j* as *N_ij_* = 1. Otherwise, *N_ij_ =* 0.
2. In the network *N,* if *N_ij_* = 1, we weigh the edge by 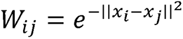 (*W* ∈ ℝ*^m×m^*). Otherwise, *W_ij_* = 0. The weighted network reflects the local structure of the *m* genes in the feature space.
3. (3) Computing the Laplacian Score of each feature. Let *y_l_* = [*y_l_*_1_,*y_l_*_2_,…,*y_lm_*]*^T^* denote the *l*-th feature values for all *m* genes. We redefine *y_l_* by removing the mean from the samples as in *Equation (4)*. The Laplacian Score (*LS*) is computed by *Equation (5)*. The lower the *LS* value, the more important the feature is. According to the scores of each feature, we finally selected the top 20 features with the smallest *LS* to be included in the prediction model for each type of cancer.

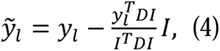

where 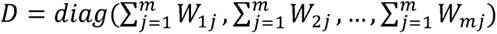, *I* = [1,1,…,1]*^T^*.

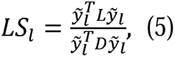

where *L = D − W*.

### Data transformation

In order to integrate multiple features of each gene into a risk score (Fig. 1), we combined the features by Hotelling and Box-Cox transformations, which converted the feature values into *p*-values. For a given scaled matrix of *m* genes with *n* features (*P* ∈ ℝ*^m×m^*, here *n* = 20), the Hotelling transformation is performed as:

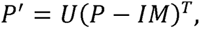

where 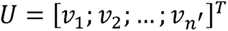 with *n′ ≤ n* as the number of chosen principal components of *P* and V = [*ν*_1_;*ν*_2_;…;*ν_n_*] are eigenvectors for the covariance matrix of *P* corresponding to decreasing eigenvalues with *λ*_1_ ≥ *λ*_2_ ≥ … ≥ *λ_n_* 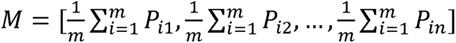·Thus, transformed *P′* ∈ ℝ*^n′×m^*. Then, the Box-Cox transformation is performed as follows,

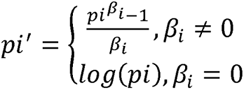

where *pi* is the *i*-th row vector of *P′* and all elements of *pi* vector are forced to be positive before being transformed, and β*_i_* is the parameter for transforming *pi* to *pi′*.

Finally, we standardized each *pi′* and calculated P-values for elements of *pi′* that is in a standard Gaussian distribution. The P-values of the elements were combined as *S*(*j*) by Fisher’s method (Fisher’s combined probability test). *S*(*j*) is termed the score of gene *j*.

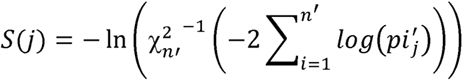

### Gibbs sampling

The scores (*S*) of genes were used as conditional probabilities in Gibbs sampling to obtain a convergent probability distribution of candidate genes. In the first round of sampling, Gibbs sampling was initiated by randomly selecting *m′* (*m′ ≤ m*) genes from the candidate genes. The *m′* genes were assumed to have equal probabilities of being selected. Then, in the second round of sampling, another set of *m′* genes was sampled from the remaining *m–m′* genes weighted by their scores (*S*). The following rounds were the same as the second round. In each round, the selected frequency 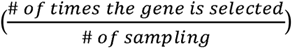 of each gene was updated. All the selected frequencies of candidate genes in the *i*-th round were denoted as a vector (*m* × 1), *Freq_i_.* When the Euclidean norm of *Freq_i_* − *Freq*_*i*−1_ was smaller than *E_Gibbs_* (*E_Gibbs_* was set as 0.01), the iteration was stopped. *Freq_last_* was assigned as the posterior probabilities (PP) of candidate genes. Then, we constructed a null distribution of PP in order to obtain the likelihood of a given gene being a cancer driver gene. The null distribution was generated by giving genes randomly weighted scores that were derived from the uniform distribution with the same range as the true scores. We generated 1,000,000 sets of null distributions of PPs for the *m* genes by running Gibbs sampling. For each gene, we obtained 1,000,000 random PPs. By counting the number of times that a random PP of a gene was larger than the real PP of the gene, an experience *p*-value was estimated. The *p*-values were adjusted by Bonferroni correction and we selected those genes with *p_adj_* < 0.01 as being significant.

### Gene set enrichment analysis

The significance of the enrichment was evaluated by the null hypothesis that the proportion of predicted cancer driver genes in a given gene set is equal to the expected proportion of the protein-coding genes in the same gene set. These human protein-coding genes were downloaded from GENCODE[38] if genes were annotated as with biotype “protein_coding”. In total, 19,350 unique human protein-coding genes were obtained. The gene set enrichment analysis was performed using a one-sided Fisher’s Exact test, and the enrichment *p*-values were corrected for multiple testing using the Bonferroni correction. We defined *p_adj_ <* 0.05 as a significant result.

### Comparison with other cancer driver prediction methods

The performance of DGAT-cancer was evaluated by the area under the precision–recall curve (AUPRC) as calculated by the perfMeas package in R[39]. The AUPRC was calculated by using known cancer driver genes collected from CGC, OncoKB and IntOGen (1,199 genes, Methods) as a positive gene set. The negative gene set contained the human genes after removing the positive genes and the genes directly interacting with the positive genes according to protein-ptotein interaction data in BIOGRID[40] (version 4.4.214, organism: human sapiens). In total, 8,153 genes were allocated to the negative gene set. The numbers of positive and negative cancer driver genes for each specific cancer are shown in Additional file 1: Table S9.

DGAT-cancer was compared to three classical methods, MutSigCV[4] (https://software.broadinstitute.org/cancer/cga/mutsig), OncodriveFML[14] (http://bbglab.irbbarcelona.org/oncodrivefml/home) and OncodriveCLUSTL[15] (http://bbglab.irbbarcelona.org/oncodriveclustl/home). MutSigCV (version 1.3.5) is a method that considers the heterogeneity of mutations as a means to identify genes that are significantly more highly mutated in cancer than expected by chance alone given background mutation processes. We ran MutSigCV using default parameters and the mutation data from Broad GDAC Firehose with the gene covariates provided by the original article[4]. OncodriveFML is a method designed to use the somatic mutation pattern across cancers to identify cancer driver genes; it was run using the online version by inputting files in “maf” format obtained from ICGC and Broad GDAC Firehose (Additional file 1: Table S1). The detailed parameters for running OncodriveFML are shown in Additional file 1: Fig. S1. OncodriveCLUSTL is a sequence-based clustering algorithm designed to detect cancer drivers. We used the online version of OncodriveCLUSTL to predict cancer driver genes based on default parameters.

### Tissue specimens and patient information

91 surgical specimens were collected from the Department of Neurosurgery, Sun Yat-sen Memorial Hospital, Sun Yat-sen University. These samples include 55 glioblastoma multiforme (GBM) specimens, 31 brain lower grade glioma (LGG) specimens and 5 non-tumor brain tissues, which had been diagnosed between 2012 and 2022. The study was approved by the Ethics Committee of Sun Yat-sen University, and informed consent was obtained from all subjects. The non-tumor brain tissues were obtained from patients with non-tumor diseases and required partial brain excision from patients with traumatic brain injury, or other diseases such as cerebral angiomas or vascular malformations.

### Experimental validation of the role of novel driver genes in cancer

In order to examine the performance of DGAT-cancer, we experimentally validated the roles of the predicted cancer drivers by using surgical specimens, a cell model and an animal model. All experimental methods are provided in the Additional file 1: Supplementary Methods.

## Results

### Application of DGAT-cancer to multiple types of cancer

DGAT-cancer was applied to the prediction of cancer driver genes in nine cancer types, Bladder urothelial carcinoma (BLCA), Breast invasive carcinoma (BRCA), Cervical and endocervical cancers (CESC), Colon adenocarcinoma (COAD), Glioblastoma multiforme (GBM), Head and neck squamous cell carcinoma (HNSC), Brain lower grade glioma (LGG), Lung adenocarcinoma (LUAD) and Stomach adenocarcinoma (STAD) whose mutation (uEMD-Mut) and gene expression (uEMD-Ex, gene expression level and MutExTDA) features were all available in TCGA database (Methods). The numbers of genes predicted to be cancer drivers by DGAT-cancer are shown in Additional file 1: Table S8.

We found significant (*p_adj_* ≤ 0.032) overlaps between the predicted cancer drivers and the cancer gene sets, CGC, OncoKB and IntOGen, respectively (Fig. 2a) compared to the background genes (19,350 protein-coding genes in ENSEMBL[41] biotype). Since cancer genes are likely to have experienced a slower evolutionary rate and stronger purifying selection than those of non-cancer, Mendelian disease, and orphan disease genes[42], we tested the enrichment of the predicted genes in the genes under selective constraint[28]. We found that the predicted cancer drivers of the nine cancer types were significantly (*p_adj_* ≤ 4.18 × 10^−4^) enriched in genes that have been under selective constraint (Fig. 2a). We further evaluated the enrichment of the predicted cancer drivers in the shRNA gene set by performing shRNA screens in 216 cancer cell lines[29] (Methods). As shown in Fig. 2a, the predicted cancer driver genes in the nine cancer types were significantly (*p_adj_* ≤ 0.034) enriched in the shRNA gene set, illustrating the potential roles of the predicted cancer drivers in cancer cell survival. The predicted cancer drivers were also enriched in cancer-related pathways defined in the Kyoto Encyclopedia of Genes and Genomes (KEGG)[43] database, e.g. Melanoma (2.90 × 10^−5^ ≤ *FDR* ≤ 0.046), Proteoglycans in cancer (5.13 × 10^−6^ ≤ *FDR* ≤ 0.033), cancer immunotherapy (*FDR* = 6.16 × 10^−3^), cell cycle (0.017 ≤ *FDR ≤* 0.024), ErbB signalling pathway (5.63 × 10^−5^ ≤ *FDR ≤* 0.024), MAPK signalling pathway (3.35 × 10^−3^ ≤ *FDR <* 0.010), Wnt signalling (1.78 × 10^−3^ ≤ *FDR ≤* 0.044), p53 signalling pathway (6.40 × 10^−3^ ≤ *FDR ≤* 0.046), and TGF-beta signalling pathway (1.61 × 10^−6^ ≤ *FDR <* 0.044) (Additional file 1: Fig. S2).

**Fig. 2.**
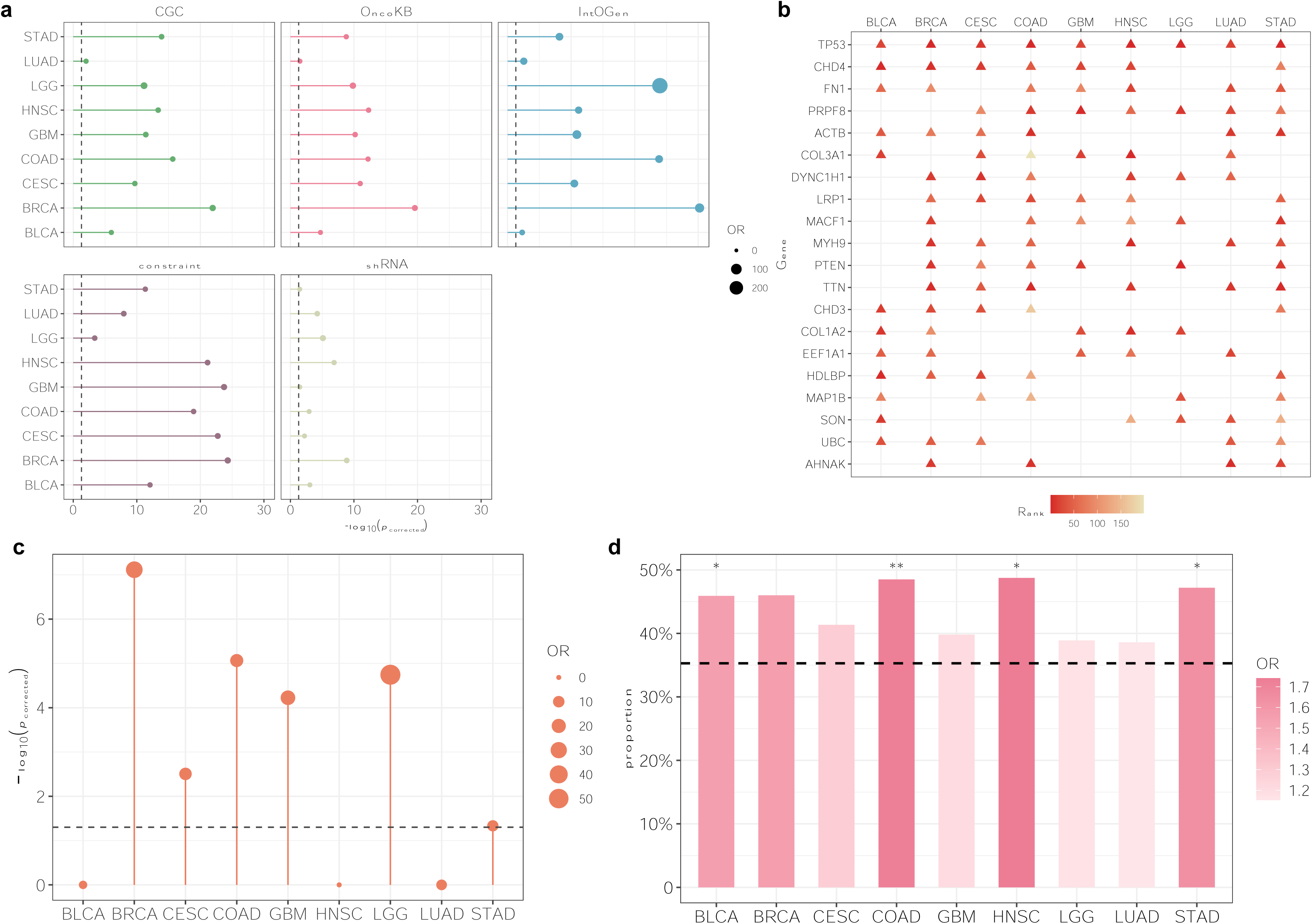
Evaluation of the cancer driver genes predicted by DGAT-cancer. **a** Enrichment of predicted cancer drivers in known cancer gene sets (CGC, OncoKB, IntOGen) and gene sets related to cancer (constraint: selective constraint genes, and genes with expression associated with functions of cancer cells). **b** Top 20 genes most frequently predicted as cancer drivers. The color depth denotes the rank order of PP (decreasing) for genes generated by DGAT-cancer in that cancer type. **c** Enrichment of predicted cancer drivers in drug-targeted genes. **d** The proportions of predicted cancer drivers whose expression levels were significantly correlated with drug activity (measured in terms of 50% growth inhibitory levels) compared to the background genes. The P-values were obtained by using Fisher’s Exact test to compare the predicted cancer drivers with predicted non-cancer drivers. *\*p_corrected_* < 0.05; *\*\*p_corrected_ <* 0.001; *\*\*\*p_corrected_ <* 0.0001.

There were 20 genes predicted as cancer drivers by DGAT-cancer in multiple cancer types (Fig. 2b). Among them, *TP53* was predicted to be a cancer driver in nine types of cancer, with predicted scores ranking between the top 1 to the top 24. The *COL1A2* gene was predicted as a cancer driver in BLCA and HNSC, with predicted scores ranking in the top 10 and the top 4, respectively. The *COL1A2* gene encodes the pro-alpha2 chain of type I collagen which is involved in the TGF-beta signalling pathway and has been reported to be associated with the migration of chondrosarcoma and fibrosarcoma cells[44]. Another gene, *PTEN,* was predicted to be a cancer driver in BRCA, CESC, COAD, GBM, LGG and STAD. *PTEN* is a tumour suppressor[45, 46] that is involved in the p53 and PI3K-Akt signalling pathways and has been found to be frequently mutated in a large number of different cancers[47, 48].

### Expression of cancer drivers predicted by DGAT-cancer correlates significantly with the level of drug activity

As shown in Fig. 2c, the predicted cancer drivers in BRCA, CESC, COAD, GBM, LGG and STAD were significantly (7.65 × 10^−8^ ≤ *p_adj_ ≤* 0.046) enriched in drug response genes. We further explored the correlation between the expression patterns of the predicted cancer driver genes and drug activities, expressed as 50% growth inhibitory levels (GI50) in the NCI-60 cell line, which were derived from CellMinerCDB[49]. The analysis was performed by calculating Pearson correlation coefficients between gene expression levels and the z-scores of negative log 10 (GI50) in all NCI-60 cell lines. We found that the expression levels of many predicted cancer drivers were significantly (*p_adj_* ≤ 0.05) correlated with drug activities (Fig. 2d, thereby illustrating the potential of these genes for clinical treatment. Briefly, the genes predicted to be cancer drivers in four out of nine cancer types (BLCA, COAD, HNSC and STAD) were significantly enriched (one-sided Fisher’s Exact test *p_adj_* < 0.036) in genes whose expression was correlated with drug activity comparing to random genes selected from the human protein gene sets.

### DGAT-cancer outperformed other methods

DGAT-cancer was compared with three other cancer driver identification methods, MutSigCV, OncodriveFML and OncodriveCLUSTL with respect to the area under the precision–recall curve (AUPRC). First, DGAT-cancer was compared with other methods of predicting cancer drivers from a set of genes (a total of 21,664 genes, Additional file 1: Table S8) having prediction scores given by at least one of the four methods. Additional file 1: Table S9 shows the numbers of positive and negative genes used in evaluating the methods. As shown in Fig. 3a, the AUPRC (in the range of 0.336 to 0.469) of DGAT-cancer in predicting cancer drivers for nine types of cancer (BLCA, BRCA, CESC, COAD, HNSC, GBM, LGG, LUAD and STAD) were the highest by comparison with the other three methods, MutSigCV (AUPRC ranged in 0.225 to 0.282), OncodriveFML (AUPRC ranged in 0.283 to 0.365) and OncodriveCLUSTL (AUPRC ranged in 0.289 to 0.356) (Fig. 3a). When assessing the AUPRC of these methods for predicting cancer drivers from genes with prediction scores provided by all four methods (Additional file 1: Table S8), DGAT-cancer also performed the best in the eight cancer types (BLCA, BRCA, CESC, COAD, HNSC, GBM, LUAD and STAD) (Additional file 1: Fig. S3a).

**Fig. 3.**
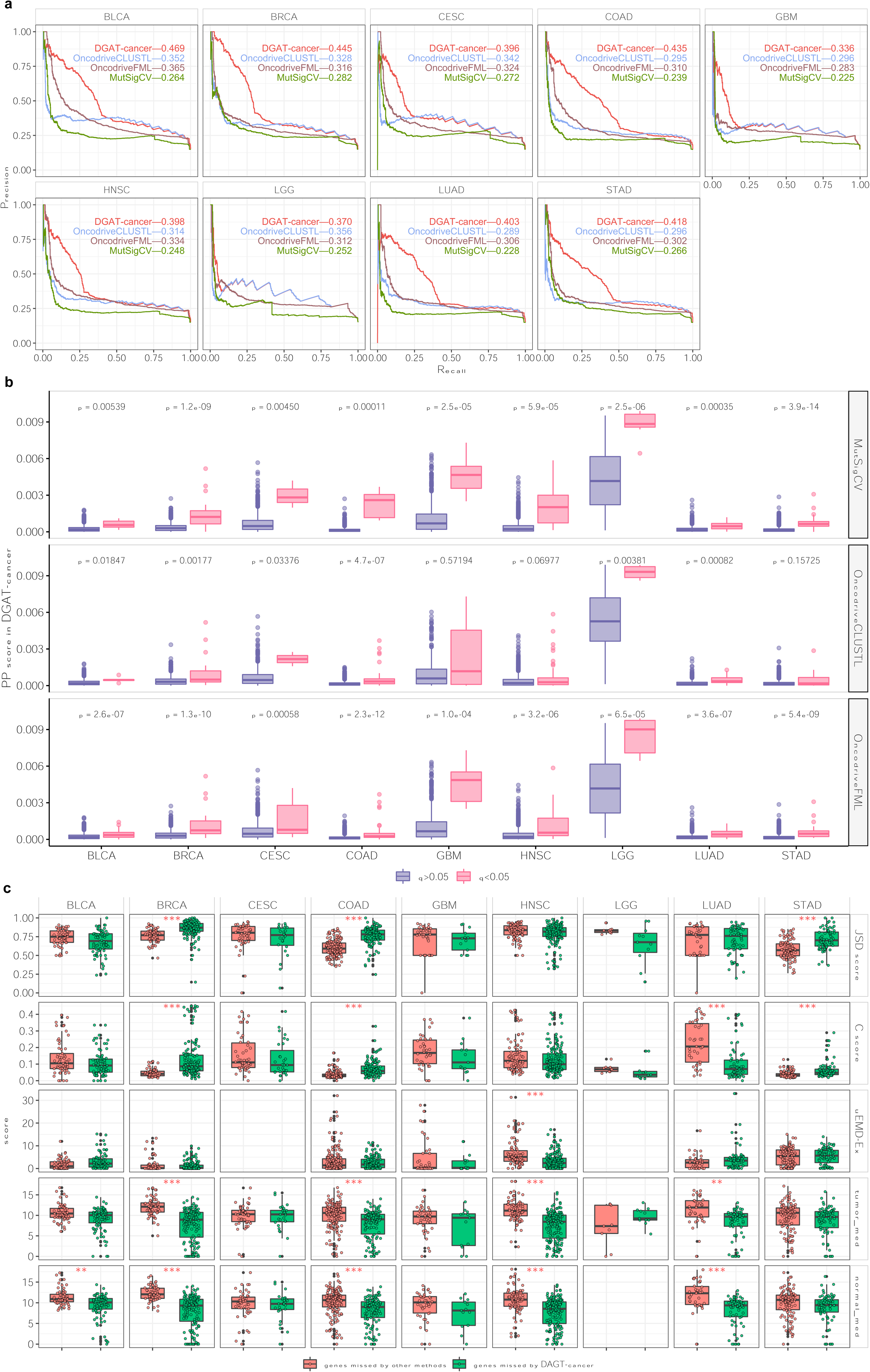
Comparison of DGAT-cancer with three methods with respect to their prediction of cancer drivers. **a** Comparison of DGAT-cancer with other methods in terms of their performance as measured by AUPRC (area under the precision–recall curve). **b** Comparing posterior probabilities (PPs) given by DGAT-cancer of genes predicted as cancer drivers by MutSigCV, OncodriveCLUSL and OncodriveFML to the genes predicted as non-cancer drivers by those methods. The P-values were obtained by means of the Wilcoxon rank-sum test. In the boxplot, the center lines represent the median, whilst the boxes represent the first and third quartiles. **c** Comparison between genes missed by DGAT-cancer and genes missed by other methods in terms of their feature scores given by DGAT-cancer. Those scores are uEMD-Ex scores, gene expression level in tumor (tumor-med), gene expression level in paracancerous tissues (normal-med), and MutExTDA scores (JSD and C score). Some boxes are blank due to the missing features in that cancer type. The difference was evaluated by Wilcoxon rank-sum test, * *p_corrected_* < 0.05; ** *p_corrected_ < **0.001***; *** *p_corrected_* < 0.0001.

Each of the four methods yielded a *p*-value to represent the probability of the predicted cancer driver being a false positive. The p-value distributions of genes generated by these four methods were then compared with those expected *p-*values from a uniform distribution using quantile-quantile plots. As shown in Additional file 1: Fig. S3b, the p-values of genes not predicted to be cancer drivers by DGAT-cancer (*p* > 0.05) in BLCA, LUAD and STAD exhibited better agreement with the expected *p*-values than those predicted by other methods. Additionally, the genes predicted (*p* < 0.05) by DGAT-cancer to be cancer drivers in all nine cancer types showed higher inflation from the expected *p*-values than the genes predicted by other methods (Additional file 1: Fig. S3c). This suggested that DGAT-cancer has enhanced potential to distinguish novel cancer drivers from random genes.

We next explored the consistency of DGAT-cancer with respect to the other methods. The posterior probabilities (PPs) given by DGAT-cancer for genes predicted to be cancer drivers (*q_risk_* < 0.05) by the other methods were compared to the PP scores of the genes not predicted to be cancer drivers *(q_risk_* ≥ 0.05) by the other methods, MutSigCV, OncodriveFML and OncodriveCLUSTL, respectively. The differences were evaluated by the Wilcoxon rank-sum test. The total number of cancer drivers with *q_risk_ <* 0.05 identified by MutSigCV, OncodriveFML and OncodriveCLUSTL are shown in Additional file 1: Table S10. As depicted in Fig. 3b, those genes predicted to be cancer drivers in BLCA, BRCA, CESC, COAD, LGG and LUAD by the three methods were given significantly higher PPs (*p* < 0.033) by DGAT-cancer than the genes not predicted to be cancer drivers by the other methods. These results suggested a high degree of consistency between DGAT-cancer and the other methods.

There were 67, 58, 50, 166, 41, 87, 9, 39 and 112 genes predicted to be cancer drivers of nine types of cancer (BLCA, BRCA, CESC, COAD, GBM, HNSC, LGG, LUAD and STAD) by DGAT-cancer, respectively but not predicted to be cancer drivers by the other three methods (OncodriveCLUSTL, OncodriveFML, MutSigCV). Moreover, 91, 152, 32, 135, 13, 177, 11, 67 and 78 genes were predicted as cancer drivers in nine types of cancer (BLCA, BRCA, CESC, COAD, GBM, HNSC, LGG, LUAD and STAD) by at least of one of the three methods, were not predicted to be cancer drivers by DGAT-cancer. We compared these genes in relation to their scores of features used in DGAT-cancer, and found that the gene expression-related features such as uEMD-Ex scores, expression values and MutExTDA scores showed a significant difference in the two groups of genes across multiple cancers (Fig. 3c and Additional file 1: Fig. S4). Specifically, the genes missed by the other three methods were expressed more highly in both tumors and paracancerous tissues than the genes that were missed by DGAT-cancer in BRCA, COAD, HNSC and LUAD. This illustrated that DGAT-cancer was more likely to detect active genes in cancer and paracancerous tissues than other methods without considering gene expression. DGAT-cancer also used features based on mutation frequency and expression profiles in tumor cohorts, JSD and C score (Methods), and fin predicting cancer drivers. For these two scores, genes missed by other methods had relatively lower JSD scores in three cancers (BRCA, COAD and STAD) and lower C scores in four cancers (BRCA, COAD, LUAD and STAD) than genes missed by DGAT-cancer (Fig. 3c), suggesting a preference for DGAT-cancer to identify genes that have highly correlated gene expression and mutation profiles.

### Gene expression-based features in tumour and paracancerous tissue served to improve DGAT-cancer

DGAT-cancer integrated both mutation-based features (uEMD-Mut) and gene expression-based features (uEMD-Ex, MutExTDA and expression values) in order to identify cancer drivers. The importance of these two types of features in the prediction was then evaluated. First, we used only uEMD-Mut to predict cancer drivers for each of the nine cancer types (Methods); this predictor was termed DGAT-Mut. The comparison between DGAT-Mut with other methods was based on a total of 23,433 genes that have the prediction score from at least one of the methods. Additional file 1: Table S11 shows the numbers of positive and negative genes used for evaluating the methods. As shown in Fig. 4, DGAT-Mut yielded higher AUPRC values (ranging from 0.272 to 0.370) for predicting cancer drivers in BLCA, BRCA, CESC, COAD, GBM, LGG and STAD than OncodriveCLUSTL (ranging from 0.288 to 0.442), OncodriveFML (ranging from 0.283 to 0.406) and MutSigCV (ranging from 0.221 to 0.282). However, the AUPRC values achieved by DGAT-Mut in predicting cancer drivers for nine cancer types were lower than with DGAT-cancer. These results illustrated the importance of integrating uEMD-Mut with the gene expression-related features for improving the prediction of cancer drivers.

**Fig. 4.**
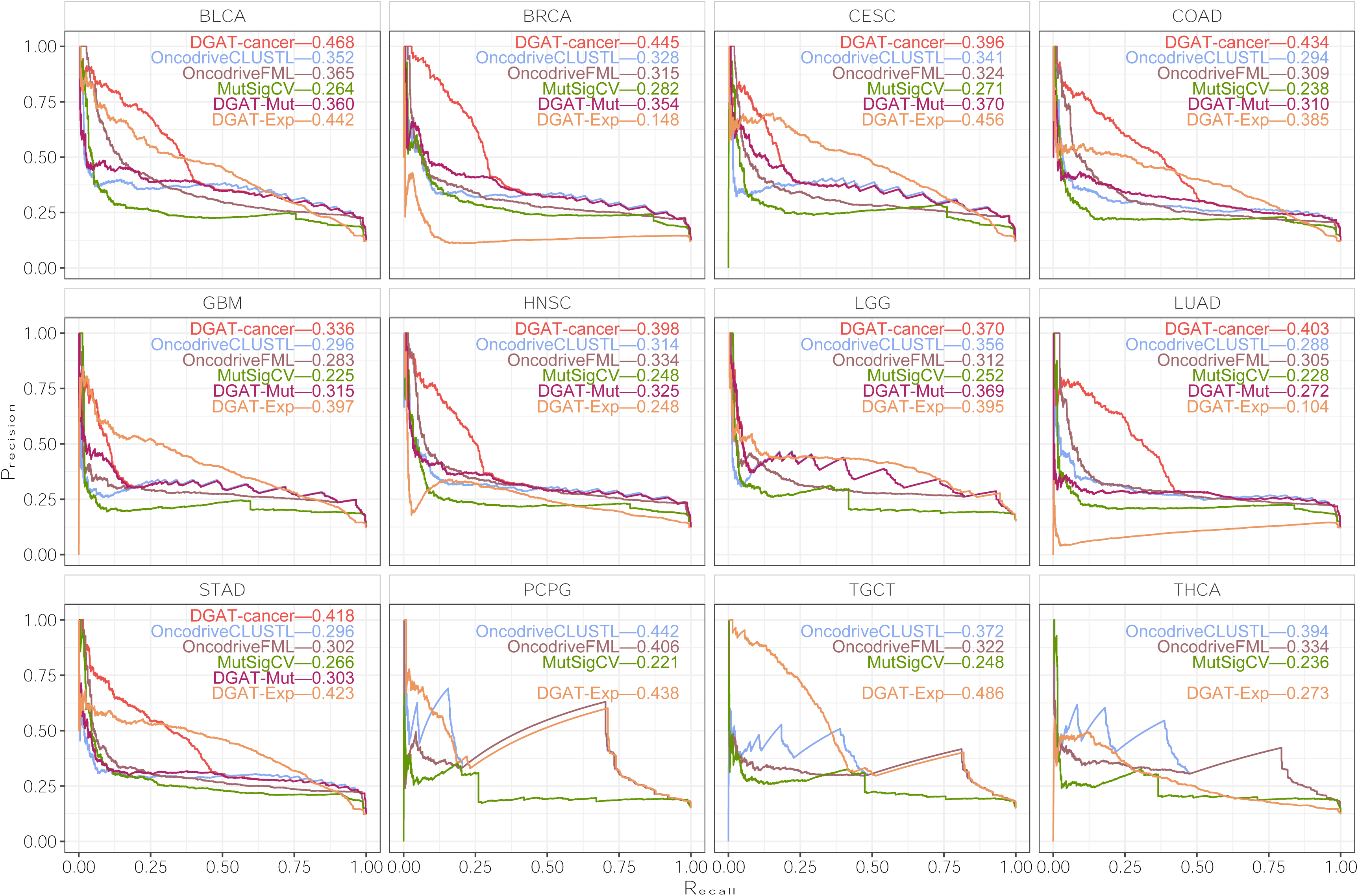
Comparing AUPRC performance of DGAT-cancer, DGAT-Mut, DGAT-Exp, MutSigCV, OncodriveFML and OcodriveCLUST to examine the impact of mutation- and gene expression-based features on improving DGAT-cancer.

When we only used gene expression-based features (uEMD-Ex, MutExTDA and gene expression values) to construct the predictive model, DGAT-Exp (Methods), it provided lower AUPRC values than DGAT-cancer in predicting cancer drivers for five out of nine types of cancer (BLCA, BRCA, COAD, HNSC and LUAD) (Fig. 4). DGAT-cancer and DGAT-Mut were not applied to the prediction of cancer drivers for three cancer types (PCPG, TGCT and THCA) because limited mutational information was available from the TCGA or ICGC databases for these three types of cancer (Methods). When DGAT-Exp was used to predict cancer drivers in these types of cancer (BLCA, CESC, COAD, GBM, LGG, STAD and TGCT), it yielded the highest AUPRC values compared to the other three methods (MutSigCV, OncodriveCLUSTL and OncodriveFML). In short, DGAT-Exp performed less well than DGAT-cancer with respect to the prediction of cancer drivers in five types of cancer (BLCA, BRCA, COAD, HNSC, and LUAD) although it yielded the best AUPRC in predicting the cancer drivers than other four methods (OncodriveCLUSTL, OncodriveFML, MutSigCV, and DGAT-Mut). This result confirms the key role of gene expression-based features in the model. Thus, combining both uEMD-Mut and expression-based features improved the performance of DGAT-cancer.

### DGAT-cancer identifies novel cancer drivers

DGAT-cancer identified many novel cancer driver genes that have not hitherto been reported to be related to cancer in CGC, OncoKB or IntOGen gene sets. Briefly, it predicted 71, 131, 60, 99, 148, 117, 20, 46 and 105 novel cancer drivers in GBM, BLCA, BRCA, CESC, COAD, HNSC, LGG, LUAD and STAD, respectively (Additional file 4: Table S12).

We next wondered if these predicted novel cancer drivers might correlate with the prognosis of the cancer patients (Methods). Relationships between the survival time of patients and gene expression were explored using a Kaplan-Meier model to analyze data from BRCA, BLCA, CESC, COAD, GBM, HNSC, LGG, LUAD and STAD in The Human Protein Atlas[33]. By comparing the number of genes whose mRNA expression level significantly (log-rank *p <* 0.05) correlated with patient survival in at least one cancer type, we observed that the predicted novel cancer drivers contain a significantly higher proportion of genes (total number 571 and proportion 96.67%) (one-sided Fisher’s Exact test, *p =* 1.03 × 10^−5^, *OR =* 2.48) that correlated with patient survival than the genes predicted to be of low probability to be cancer drivers (removing genes contained in CGC, OncoKB, IntOGen and predicted cancer drivers, total number 7,723 and proportion 92.14%). For example, *CD44,* a gene that was predicted to be a novel cancer driver in LUAD (with scores ranked in the top 48 and *p_adj_ <* 10^−6^) was found to be significantly correlated with patient survival for COAD (*p* = 5.00 × 10^−2^), GBM (*p* = 3.50 × 10^−3^), LGG (*p* = 3.50 × 10^−3^)and HNSC (*p* = 1.46 × 10^−2^). It has been reported that the expression level of *CD44* exhibits a positive correlation with PD-L1 protein in LUAD patients [52]. *TTN* was another novel gene predicted to be a cancer driver in six types of cancer, BRCA, CESC, COAD, HNSC, LUAD and STAD (with scores ranked in the top 2 to top 39, and *p_adj_ <* 10^−6^) and the expression level was found to correlate with survival in BRCA (*p* = 1.00 × 10^−2^), BLCA (*p* = 1.34 × 10^−2^), GBM (*p* = 3.71 × 10^−2^), LGG (*p* = 3.71 × 10^−2^) and HNSC (*p* = 1.49 × 10^−2^). Finally, *EEF1A1* was predicted to be a cancer driver in BLCA, BRCA, GBM, HNSC and LUAD (ranked 21 to ~68, *p_adj_ <* 10^−6^). Further analysis indicated that the expression of *EEF1A1* was significantly correlated with the survival of patients in CESC (*p* = 7.86 × 10^−3^), COAD (*p* = 2.53 × 10^−2^), GBM (*p* = 1.39 × 10^−2^) and LGG (*p* = 1.39 × 10^−2^). A previous study has indicated that *EEF1A1* is expressed at a significantly higher level in glioblastomas than in non-neoplastic white matter[53]. However, no studies have so far been performed to establish a relationship between *EEF1A1* and GBM. We, therefore, performed further experiments to validate a possible role for *EEF1A1* in glioma.

## Experimental validation of a role for EEF1A1 in glioma

### *EEF1A1* expression is increased in glioma and correlates with a poor prognosis

We compared the mRNA expression levels of *EEF1A1* in 698 glioma samples from the TCGA GBM/LGG dataset and 1,157 normal brain samples from GTEx[50]. The results showed that *EEF1A1* expression was upregulated in tumor tissue relative to normal brain tissue, as well as significantly increased in GBM relative to LGG (Both P<0.001, Fig. 5a). Moreover, the expression level of *EEF1A1* in the glioma samples from TCGA was found to negatively correlate with the overall survival of patients (P=0.026, Fig. 5b).

**Fig. 5.**
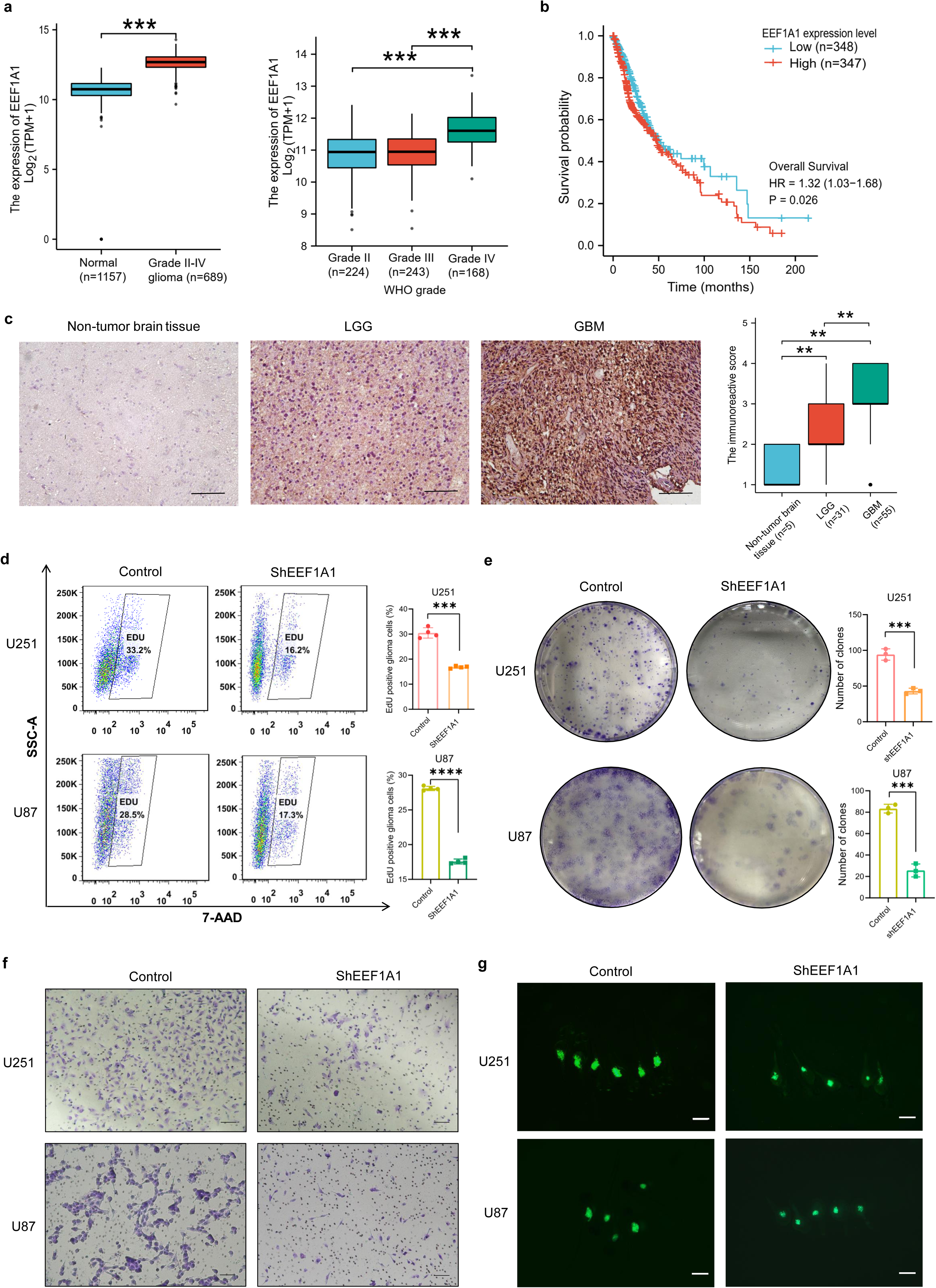
*EEF1A1* is highly expressed in glioma and plays a role in regulating the proliferation and migration of glioma cells. **a** Profile of *EEF1A1* mRNA expression in normal, LGG (WHO grade II–III glioma), or GBM (WHO grade IV glioma) patients in the GTEx and TCGA datasets. *EEF1A1* expression was significantly upregulated in glioma relative to normal brain tissue (left) (Mann-Whitney U test), as well as significantly increased in GBM relative to LGG (right) (Kruskal-Wallis test, adjusted by Bonferroni correction). **b** Kaplan–Meier overall survival plot showing that the survival rate of glioma patients with high *EEF1A1* expression (red) in the TCGA data set was significantly lower than those with low *EEF1A1* expression (blue) (two-sided log-rank test). **c** *EEF1A1* protein expression in non-tumor brain tissue, LGG and GBM samples using immunohistochemical analysis. Non-tumor brain tissues were obtained from patients with non-tumor brain diseases who had undergone surgical resection. Representative images of immunohistochemical analysis. Scale bar=10μm. The summary of *EEF1A1* protein expression profile in C (Mann-Whitney U test) was on the right. **d** EdU flow cytometry assay showed that the cell proliferation rate was significantly decreased in U251 and U87 cells with the knockdown of *EEF1A1*. **e** *In vitro* colony formation of U251 and U87 cells was decreased after *EEF1A1* knockdown compared to the control. **f** Representative images of the transwell migration assay for migrated cells stained with crystal violet in control cells and shEEF1A1 knockdown U251 and U87 cells. Scale bar = 1mm. **g** Representative images of the zebrafish xenograft model used to analyze the proliferation of U251 cells after *EEF1A1* knockdown by measuring GFP fluorescent foci. Scale bar = 0.5mm. Unpaired two-sided t test. \**p* < 0.05; \*\**p* < 0.01; \*\*\**p* < 0.001; \*\*\*\**p* < 0.0001.

The *EEF1A1* protein level was examined in 91 glioma specimens by immunohistochemical analysis; these included 55 GBM tissues, 31 LGG tissues and 5 non-tumor brain tissues. Relative to non-tumor brain tissues, LGG exhibited a significantly higher *EEF1A1* level in LGG (*p* = 6.59 × 10^−3^, Fig. 5c) and GBM (*p =* 2.12 × 10^−3^, Fig. 5c). Notably, the expression of *EEF1A1* protein was significantly increased in GBM as compared with LGG (*p* = 4.02 × 10^−3^, Fig. 5c). When we cultured GBM cell lines (U251 and U87 cells) and LGG cell lines (Hs683), we found that *EEF1A1* protein was more highly expressed in GBM cells (U251 and U87) than in LGG cells (Hs683) (*p* = 1.01 × 10^−3^ for U87 *vs*. Hs683, *p* = 6.82 × 10^−4^ for U251 *vs*. Hs683, Additional file 1: Fig. S5). Thus, we surmised that *EEF1A1* may be involved in glioma tumorigenesis.

### Knockdown of *EEF1A1* inhibited the proliferation and migration of glioma cells

To examine the role of *EEF1A1* in glioma tumorigenesis, we used shRNA to knock down *EEF1A1* expression in U251 and U87 cells. To assess knockdown efficiency, we performed real-time quantitative PCR (RT-qPCR) and Western blot assays. The results showed that compared with the control group, *EEF1A1* mRNA in the knockdown group decreased by about 77% and 86% in U87 and U251 glioma cells (*p* = 2.82 × 10^−17^ for U87 and *p* = 8.35 × 10^−11^ for U251, Additional file 1: Fig. S6a), respectively. Western blot experiments showed that *EEF1A1* protein expression in the knockdown groups decreased by 65% and 45% in U87 and U251 glioma cells (*p* = 1.67 × 10^−2^ for U87 and *p* = 2.10× 10^−3^ for U251), respectively (Additional file 1: Fig. S6b).

To assess the effect of *EEF1A1* knockdown on the proliferation of U251 and U87 cells, we performed a Cell Counting Kit-8 (CCK-8) assay, an EdU flow cytometry assay and a colony formation assay. The results showed that U251 and U87 cells with reduced *EEF1A1* expression exhibited a significant decrease in cell viability within 72h according to the CCK-8 assay (−25% and *p =* 9.19 × 10^−4^ for U251, −11% and *p* = 1.49 × 10^−2^ for U87, Additional file 1: Fig. S6c), whilst the proportions of EdU-positive cells were decreased by 45% for U251 *(p =* 1.32 × 10^−5^) and by 38% for U87 (*p* = 7.74 × 10^−9^) (Fig. 5d), with the numbers of colonies being decreased by 54% for U251 (*p* = 5.80 × 10^−4^) and by 69% for U87 (*p* = 1.61 × 10^−4^) (Fig. 5e) compared with the control group. These results demonstrated that *EEF1A1* knockdown significantly inhibited the proliferation of U251 and U87 cells.

The migration capacity of glioma cells also decreased significantly after *EEF1A1* knockdown. We found that after *EEF1A1* knockdown, the percentage of cells migrating through the transwell plate significantly decreased (−69% and *p* = 4.31 × 10^−11^ for U251, −71% and *p* = 3.96 × 10^−10^ for U87, Fig. 5f and Additional file 1: Fig. S6d). A scratch-wound healing assay yielded a similar result. A significantly shorter migration distance was observed in U251 cells with *EEF1A1* knockdown (−44%, *p* = 2.62 × 10^−2^, Additional file 1: Fig. S6e) and U87 cells (−30%, *p* = 2.97 × 10^−3^, Additional file 1: Fig. S6f). To assess the proliferation of glioma cells *in vivo*, U251 and U87 cells transfected with *EEF1A1* shRNA or control shRNA were injected into zebrafish and the areas of GFP fluorescent foci were measured. As shown in Fig. 5g, U251 and U87 cells with *EEF1A1* knockdown exhibited a significant decrease in the areas of fluorescent foci (−50% and *p* = 1.96 × 10^−3^ for U251, −41% and *p* = 3.20 × 10^−2^ for U87, Additional file 1: Fig. S6g). Taken together, these results suggest that *EEF1A1* plays a role in regulating the proliferation and migration of glioma cells.

### Knockdown of *EEF1A1* increased temozolomide (TMZ) sensitivity in glioma cells

In order to explore the role of *EEF1A1* in the sensitivity of glioma cells to temozolomide (TMZ), U251 and U87 cells with *EEF1A1* knockdown, and control glioma cells were cultured in different concentrations of TMZ. The results showed that knockdown of *EEF1A1* significantly decreased cell viability at different concentrations of TMZ (*p* = 2.66 × 10^−2^ for U251 cells and *p* = 9.10 × 10^−3^ for U87 cells, Fig. 6a), as well as at the half maximal inhibitory concentration (IC50) of TMZ in glioma cells (−41% and *p* = 2.94 × 10^−3^ for U251, −44% and *p* = 6.44 × 10^−3^ for U87, Fig. 6b). Next, a colony formation assay was performed in cells with treatment of TMZ (50μM). In the presence of TMZ, knockdown of *EEF1A1* significantly reduced colony formation (−64% and *p* = 2.93 × 10^−5^ for U251 cells, −58% and *p* = 1.60 × 10^−4^ for U87 cells, Fig. 6c and d).

**Fig. 6.**
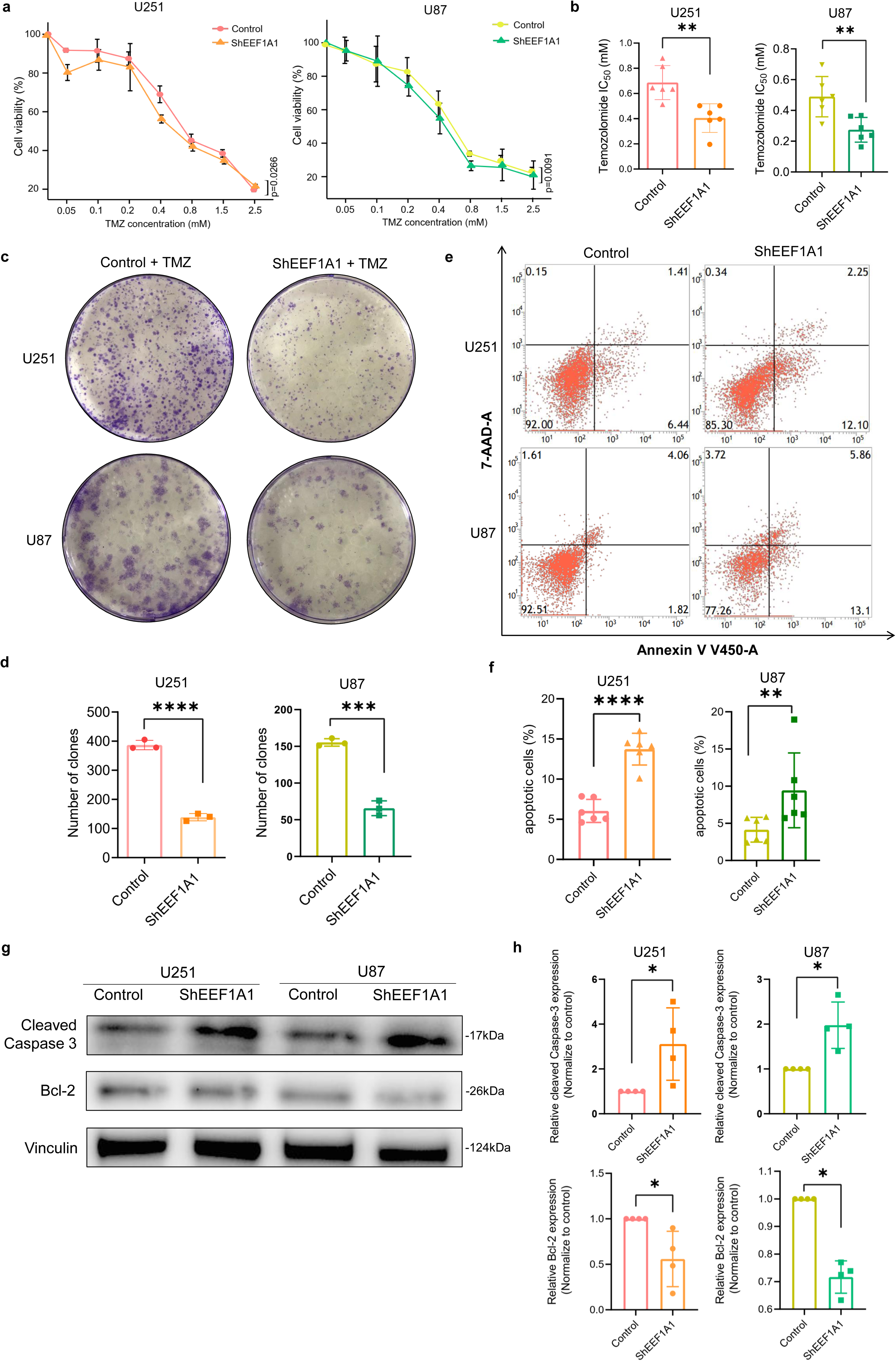
Knockdown of *EEF1A1* improved TMZ sensitivity in glioma cells. **a** Knockdown of *EEF1A1* significantly decreased the viability of U251 and U87 cells under different concentrations of TMZ (24h for U251 cells and 48h for U87 cells) (*n*=6, paired two-sided t test). **b** Knockdown of *EEF1A1* significantly decreased the IC50 of TMZ in U251 and U87 cells (*n*=6, unpaired two-sided t test). **c** In the presence of TMZ (50 ug/mL), the knockdown of *EEF1A1* significantly reduced colony formation. **d** Quantitation of colony formation in (**c**) (*n*=3, unpaired two-sided t test). **e** Flow cytometry results showing the proportions of apoptotic cells (the Annexin V V450-positive cells) in the EEF1A1-knockdown and control groups in U251 and U87 cells. **f** The summary of six independent experiments of (**e**) (unpaired two-sided t test). **g** Western-blot showing the cleaved caspase-3 and bcl-2 protein expression in the *EEF1A1*-knockdown and control groups in U251 and U87 cells. **h** Quantitation of four independent experiments of (**g**). (Mann-Whitney U test) *\*p* < 0.05; *\*\*p* < 0.01; ***p < 0.001; ****p < 0.0001.

Culturing U251 and U87 cells in each group with 200μM TMZ, we observed that the cell viability of the *EEF1A1*-knockdown group was significantly lower (*p* < 0.01) than that of the control group according to CCK-8 assays (Additional file 1: Fig. S7a). These results suggest that the knockdown of *EEF1A1* is able to inhibit the proliferation of glioma cell lines in the presence of TMZ.

To investigate the effect of *EEF1A1* on the apoptosis of glioma cells treated by TMZ, we cultured U251 and U87 cells with 200μM TMZ for 24h, and assessed cellular apoptosis by means of a flow cytometer. The results showed that the proportion of apoptotic cells in the *EEF1A1*-knockdown group was significantly higher than that in the control group (+127% and *p* = 1.60 × 10^−5^ for U251, +129% and *p* = 6.50 × 10^−3^ for U87, Fig. 6e and f). Moreover, apoptosis-related proteins also showed significant changes after the knockdown of *EEF1A1*. Thus, cleaved caspase-3, the effector of apoptotic activity, was significantly increased, whilst bcl-2, an inhibitor of apoptosis, was significantly decreased in *EEF1A1*-knockdown glioma cells (Both *p* = 2.86 × 10^−2^) (Fig. 6g and h). These findings indicated that the decreased expression of *EEF1A1* can promote apoptosis in glioma cells, thereby increasing the TMZ sensitivity of glioma cells.

## Discussion

The computational identification of cancer driver genes is key to improving our understanding of the underlying mechanisms of tumorigenesis. Most of the current methods for the identification of cancer drivers rely on the use of a single type of genomic data, such as mutations or gene expression[51]. Recently, approaches have been developed that integrate somatic mutation rates with biological networks for the prediction of cancer driver genes[52]. However, none of these approaches have considered the functional impact of mutations and tissue type-specific gene expression networks, potentially resulting in the introduction of false positives in the prediction. Here, we have developed a method, DGAT-cancer, which integrates the predicted pathogenicity of mutation profiles from cancer cohorts and a healthy population with the gene expression profiles of tumors and paracancerous tissues into a risk score that is capable of predicting cancer drivers. The method takes advantage of the huge amount of mutation data from individual samples generated by the 1000 Genomes Project, the TCGA project and the ICGC project to obtain the distribution differences of the predicted pathogenic scores of mutations in the healthy population and in tumor tissues. These differences were then integrated with the topological network of gene expression in tumor tissues and gene expression data from paracancerous tissues. To filter out the non-redundant features, we selected those features that allowed the preservation of the local geometric structure of the feature space by an unsupervised method, the Laplacian Score. DGAT-cancer is capable of detecting cancer drivers for any type of cancer by inputting mutation and/or gene expression data.

Comparing DGAT-cancer with three existing methods, it achieved the highest AUPRC in predicting cancer drivers from five out of nine types of cancer while OncodriveFML achieved the highest AUPRC in the prediction of cancer drivers for four types of cancer. The reliability of DGAT-cancer was further evaluated by comparing the *p*-values of the predicted cancer drivers to the expected *p*-values. The result showed that DGAT-cancer is a powerful tool for discriminating cancer drivers from random genes. Compared to other methods, DGAT-cancer has the ability to recognize genes with a higher expression level in tumors and paracancerous tissues (Fig. 3c), suggesting the importance of introducing tissue-specific gene expression as a feature in predicting cancer driver genes. Moreover, for the cancer types with larger sample numbers, such as BLCA, CESC, HNSC and STAD, DGAT-cancer exhibits a preference for the prediction of genes with higher uEMD-mut scores calculated using the information on tumor mutations and germline variants in the healthy population (Additional file 1: Fig. S4). Additional analysis indicated that employing only mutational information in cancer driver prediction was insufficient to achieve enhanced performance. As indicated in Fig. 4, DGAT-Mut only performed better than DGAT-Exp in its prediction of cancer drivers in BRCA, HNSC and LUAD. One likely reason is that the accuracy of DGAT-mut is influenced by the 19 features for the prediction of the pathogenicity of mutations. Especially, DGAT-cancer performed the best compared to all other methods in predicting cancer drivers for three types of cancers, LGG, PCGC and TGCT which are cancer types lacking gene expression data in paracancerous tissues (Fig. 4). This result may reflect the important roles of gene expression data in tumor tissue in predicting cancer drivers.

DGAT-cancer predicted many novel candidate genes that require further experimental validation. Thus, *AHNAK* was predicted as a cancer driver in BRCA, COAD, LUAD and STAD. It encodes a large structural scaffold protein and has been reported as a tumour suppressor in the proliferation and invasion of triple-negative BRCA[53] and LUAD[54]. *AHNAK* may function as a tumour suppressor through the inhibition of p-ERK and ROCK1 in COAD[55]. *DST* was predicted as a cancer driver in BRCA, COAD, GBM and STAD. Up-regulation of gene *DST* has been observed in the ductal carcinoma *in situ* component of BRCA[56]. *MUC5B* was predicted as a cancer driver in BRCA, COAD and STAD. Abnormal expression of *MUC5B* in COAD has been found to be associated with low expression of p53[57]. *COL1A2* was predicted to be a cancer driver in BLCA, BRCA, GBM, HNSC and LGG. *COL1A2* has been found to be associated with tumorigenesis in HNSC[58], and is a hub gene in the perineural invasion (PNI)-associated co-expression module, where PNI is a key pathological feature of HNSC[59]. *KRT14* was predicted to be a cancer driver in HNSC. It has been reported to be upregulated in HNSC[60] and its expression level in HNSC tumours is associated with patient survival[61]. *LOXL2* was predicted to be a cancer driver of HNSC. A previous study has shown that a novel splice variant in *LOXL2* upregulated *LOXL2* in HPV-negative HNSC and enhanced proliferation, migration, and invasion in HPV-negative HNSC cells[62]. The silencing of *LOXL2* inhibited cell migration and invasion in HNSC cell lines[63].

This is the first study to predict *EEF1A1* to be a cancer driver in GBM. *EEF1A1* protein has been reported to interact with key components of the pathway that regulates the synthesis of apoptosis-related proteins[64–66]. Additional roles for *EEF1A1* in apoptosis have been studied in gastric cancer[67], ovarian cancer[68], renal cell carcinoma[69] and lung cancer[70]. Here, we found that the knockdown of *EEF1A1* inhibits apoptosis in glioma cells when the cells are treated with TMZ. The underlying mechanisms require further investigation. However, when we explore the role of *EEF1A1* in glioma cells without TMZ, we did not observe any change in apoptosis when *EEF1A1* was knocked down (Additional file 1: Fig. S7b). Further experiments are required to establish the precise nature of the role that *EEF1A1* plays in glioma tumorigenesis.

There are many ways to further improve the accuracy of DGAT-cancer. First, DGAT-cancer is dependent on mutation data from the general population and cancer population. When the sample size is increased, more mutational information will become available which will help to improve the accuracy of predictions. Moreover, DGAT-cancer only takes account of mutation and gene expression data in its predictions. Other features, such as the protein or RNA structures around the mutations could also contribute important information that might be of use in discriminating cancer drivers. Finally, since the performance of DGAT-cancer is influenced by pathogenicity predictions for mutations, more accurate predictors for determining the pathogenicity of mutations could help to improve DGAT-cancer.

## Conclusions

We demonstrate that DGAT-cancer is powerful in predicting cancer drivers using mutation and/or gene expression data, and has a superior performance compared to three commonly used methods. The importance of gene expression data and mutation information in predicting cancer drivers was evidenced. DGAT-cancer has broadened our path to detect cancer driver genes and shed a light on cancer therapy.

## Declarations

### Ethics approval and consent to participate

Not applicable.

### Consent for publication

Not applicable.

### Availability of data and materials

The DGAT-cancer method is available as an open-source software package on the GitHub repository (https://github.com/Dan-He/DGAT-cancer). Simple somatic mutation (SSM) data from 12 cancer cohorts were downloaded from the Broad Institute GDAC Firehose Portal (http://gdac.broadinstitute.org/), the International Cancer Genome Consortium (ICGC) Data Portal (https://dcc.icgc.org/releases/current/Projects/) and The Cancer Genome Atlas (TCGA) (https://www.cancer.gov/about-nci/organization/ccg/research/structural-genomics/tcga). The germline mutation data of a healthy population were collected from Phase 3 of the 1000 Genomes Project (https://www.internationalgenome.org/data, GRCh38). RNA-seq data of tumors from 12 cancer types in TCGA were downloaded from the UCSC Xena platform (http://xena.ucsc.edu/).

Additional information is available at the website.

### Competing interests

All authors declared that they have no competing interests.

## Funding

This work was supported by the National Key Research and Development Program of China (2020YFB0204803), the Natural Science Foundation of China (81801132, 81971190, 61772566), Guangdong Key Field Research and Development Plan (2019B020228001, 2018B010109006, and 2021A1515010256), Introducing Innovative, Guangzhou Science and Technology Research Plan (202007030010).

## Author’s contributions

HY.Z designed the study. D.H performed the model construction and analyses, with assistance from Z.L and T.L. L.L performed the experiments, with assistance from HS.Z, F.L, S.L and B.L. D.H, L.L and Z.L wrote the manuscript. HY.Z and D.N.C supervised the study. All authors discussed the results and interpretation and contributed to the final version of the paper.

## Acknowledgments

The authors thank many resources for making data available.

## Notes

### Competing Interest Statement

The authors have declared no competing interest.

