## Additional file 1 for "Integrating gene mutation spectra from tumors and the general population with gene expression topological networks to identify novel cancer driver genes"

### Integrating spectrum of gene mutations in tumor and the healthy population with gene expression topological networks to identify cancer driver genes

**This document contains Supplementary Methods, Tables and Figures.**

### Supplementary Methods

#### Cell culture

Human U251 and U87 glioblastoma cells were purchased from Procell Life Science & Technology (Wuhan, China). Cells were cultured in DMEM medium (Gibco, 11965092) with 10% fetal bovine serum (FBS) and 1% Penicillin-Streptomycin at 37℃ with 5% CO_2_.

#### Generation of stable EEF1A1 knockdown cell lines

Lentiviral vectors containing EEF1A1 short hairpin RNA (shRNA) or negative-control shRNA were obtained from Kidan Biosciences (Guangzhou, China). The siRNA target sequence used in the EEF1A1 knockdown group (shEEF1A1 group) was 5’-CCTCTCCAGGATGTCTACAAA-3’. The scrambled siRNA sequence used in the control group was 5’-TTCTCCGAACGTGTCACGTT-3’. For cell transfection, cells seeded in six-well plates (3x10^4^ cells/well) were transduced with the constructed lentiviral shRNA particles (Lv-shEEF1A1 or Lv-scramble) in a complete medium with 5ug/mL polybrene. The medium was replaced with fresh medium 24h after infection. 72h after transfection, the tumor cells were stably infected with viral particles, and cells that expressed green fluorescent protein (GFP) were isolated by selection in the presence of 2μg/mL puromycin.

#### CCK-8 assay

For analysis of cell proliferation, 1×10^4^ cells were plated into 96-well plates and cultured. After cell adhesion, cells were cultured for 0, 24, 48 and 72h, respectively. At each time point, CCK-8 solution (10%) was added to each well. Cells were incubated at 37℃ for 2h. Thereafter, the absorbance at 450nm wavelength was measured according to the manufacturer’s instructions.

For detection of the half maximal inhibitory concentration (IC50) of TMZ, 1×10^4^ cells were placed into 96-well plates and exposed to different doses (0, 0.05mM, 0.1mM, 0.2mM, 0.4mM, 0.8mM, 1.5mM, 2.5mM) of TMZ for 24 or 48h. At each time point, 10μl CCK-8 was added into each well, and cells were cultured for 2h. The optical density (OD) value was measured at 450nm using a microplate reader (Bio-Rad, USA). Cell viability was normalized to the control group (0μg/mL TMZ), and the IC50 of TMZ was determined according to the viability curve.

#### Flow cytometry analyses

For analysis of cell proliferation, EdU Flow Cytometry Assay kits (Cy5) (APExBIO, #K1078) were used to detect cell proliferation according to the manufacturer's instructions. Briefly, cells were pre-incubated for 2h with EdU to allow its incorporation into DNA. Then the cells were fixed with 4% paraformaldehyde for 15min, permeabilized with 0.5% Triton X-100 in PBS for 20min, and incubated with 100μL Click-iT reaction mix for 30min. Cells were then analyzed by flow cytometry (BD FACS Calibur flow cytometer) to determine the proportion of EdU-positive cells, and the data were analyzed by FlowJo software.

The Annexin V-EV450/7-AAD Apoptosis Kit (Elabscience, E-CK-A234) was used to measure cell apoptosis. U251 and U87 cells were treated with 200μM TMZ for 24h before being stained according to the instructions of the antibody manufacturers. Briefly, we collected the cells, washed them with cold PBS, and resuspended them in 100μl binding buffer. We then added 2.5μl Annexin V-V450 and 7-AAD, and incubated them in the dark for 15min. Subsequently, another 100μl binding buffer was added and flow cytometry performed to determine the percentage of V450-positive and/or 7-AAD-positive cells. Cells at the early stage of apoptosis were identified as V450-positive and 7-AAD-negative, whereas cells at the late stage of apoptosis or already dead were identified as V450-positive and 7-AAD-positive.

#### Wound healing assay

Cells were seeded in 6-well plates at a density of 5x10^5^ cells per well. After 24h, the culture medium was removed and serum-free medium was added. A straight wound was scratched using a 10μl pipette tip. The cells would migrate through the scratch and heal the wound. Photographs were taken using an inverted microscope at 0, 12, 24, 48 and 72h. The results at 48h for U251 cells and 24h for U87 cells are shown.

#### Colony formation assay

For the analysis of cell proliferation, 1000 cells were seeded in 6-well plates per well. The culture medium was replaced every 3 days. Colonies were fixed with 4% paraformaldehyde and stained with 0.5% crystal violet for 20min after 10 days of culture.

For the analysis of TMZ resistance, 2000 cells were seeded in 6-well plates per well. TMZ was added to the culture medium at a concentration of 50μM. The culture medium was replaced every 3 days. Colonies were fixed with 4% paraformaldehyde and stained with 0.5% crystal violet for 20min after 14 days of culture.

#### Transwell assay

Cells (2x10^5^ cells/well) were cultured in a transwell chamber (Corning, #3422-1) containing FBS-free medium in the upper chamber and 10% FBS in the lower chamber at 37℃ with 5% CO_2_. After 24h, 4% paraformaldehyde was used to fix the cells that had migrated to the lower chamber, and 0.5% crystal violet was used to stain the cells for 20 min. The cells were photographed and counted under a microscope.

#### RNA extraction and quantitative real-time PCR

TRIzol reagent (Invitrogen, #15596-026) was used to extract total RNA from cells, which was then reverse transcribed with Vazyme HiScript III RT SuperMix (#R323-01). Quantitative real-time PCR (qPCR) was performed on an ABI Prism 7500 Sequence Detection System using ChamQ Universal SYBR qPCR Master Mix (Vazyme, #Q711-02). The RNA primer sequence used in the experiments: EEF1A1: Forward: 5’-TGTCGTCATTGGACACGTAGA-3’, Reverse: 5’-ACGCTCAGCTTTCAGTTTATCC-3’; GAPDH: Forward: 5’-GGAGCGAGATCCCTCCAAAAT-3’, Reverse: 5’-GGCTGTTGTCATACTTCTCATGG-3’. The relative expression of the genes of interest was calculated using the 2^-ΔΔt^ method.

#### Western blotting

Glioma cells were lysed with cell lysis buffer containing 0.1% protease inhibitor, 0.1% phosphatase inhibitors and 1% PMSF. The supernatant fractions (protein samples) were collected after centrifugation at 13,000 rpm for 15min at 4℃. After separation on 8% SDS-PAGE, protein samples were transferred to nitrocellulose membranes prior to blotting with 5% BSA for 60min at room temperature. The membranes were then incubated overnight at 4℃ with the following antibodies: anti-EEF1A1 (Huabio, #ET7107-75), anti-GAPDH (Huabio, #EM1901-57), anti-SOX2 (Huabio, #EM1708-84), anti-Bcl-2 (Abclonal, A0208), anti-Caspase3 (Huabio, #ET1602-47). Appropriate secondary antibodies were incubated for 2h at room temperature. Quantification was performed with ImageJ software.

#### Immunohistochemical (IHC) staining

Immunohistochemical (IHC) staining was performed according to the instructions of the antibody manufacturers. Briefly, tissue slides or tissue microarray (TMAs) of human surgical specimens were deparaffinized, rehydrated through an ethanol series, followed by antigen retrieval with sodium citrate solution. The sections were then incubated with 1% Triton-X100 for 15min, 3% H_2_O_2_ for 10min, blocked with 5% normal goat serum for 30min at room temperature, and then incubated with appropriate primary antibodies at 4°C overnight. IHC staining was performed with horseradish peroxidase (HRP) conjugates employing diaminobenzine (DAB) detection. The immunoreactive score (IRS) was based in part on the proportion of positive cells (A): 0 = no positive cells; 0.3 = 30% positive cells, 0.5 = 50% positive cells, 0.8 = 80% positive cells, 1 = 100% positive cells, and in part the intensity of staining (B): 1 = no color reaction; 2 = mild reaction; 3 = moderate reaction; 4 = intense reaction. The immunoreactive score (IRS Score) was the product of A and B (i.e. A×B).

#### Zebrafish xenograft model

Zebrafish fertilized eggs (Zebrafish Technology Platform, Sun Yat-sen University) were randomly divided into four groups (n=50 per group) and placed on a 10cm petri dish coated with 3% agarose. U251 or U87 cells transfected with lentiviral vectors containing EEF1A1 shRNA or control scrambled shRNA were manually counted by a Picospritzer microinjection device (Parker Inc), and the cells were then injected into the yolk sac of zebrafish embryos (approximately 200-300 cells per egg) in groups. After injection, zebrafish eggs were incubated at 33℃ for 72h. The zebrafish were then anesthetized with 0.02% tricaine (Sigma) and observed under fluorescence microscopy. Zebrafish with no fluorescent foci were excluded for further analysis. For the experimental procedure, three different investigators were involved as follows: the first investigator provided the transfected cells and only they were aware of the treatment assignment. A second investigator was responsible for the injection process, whilst the third monitored the fluorescence in the yolk sac. The experiments were performed in accordance with the national or institutional guidelines for animal care and use in China.

#### Statistical analysis

All results were analyzed using GraphPad Prism software (Version 7.0). Data were analyzed by means of an unpaired two-sided t-test (data with normal distribution) or a Mann-Whitney t-test (data with non-normal distribution). All data were expressed as the mean ± standard error of the mean (SEM) and were obtained from at least three independent experiments. A difference of p<0.05 was considered to be statistically significant (*$p<0.05$, **$p<0.01$, ***$p<0.001$, ****$p<0.0001$).

#### Supplementary Tables

**Table S1. Data sources for downloading somatic mutations.**

| **Cohort** | **No. of samples** | **No. of genes** | **Source** |
| --- | --- | --- | --- |
| BLCA | 411 | 17368 | ICGC |
| BRCA | 1020 | 16942 | ICGC |
| CESC | 289 | 16313 | ICGC |
| COAD | 402 | 18093 | ICGC |
| GBM | 388 | 15332 | ICGC |
| HNSC | 508 | 16613 | ICGC |
| LGG | 508 | 12915 | ICGC |
| LUAD | 516 | 17404 | ICGC |
| STAD | 439 | 17721 | ICGC |
| THCA | 492 | 6462 | ICGC |
| PCPG | 179 | 2462 | firehose |
| TGCT | 149 | 6778 | firehose |
| 1000G | 2548 |  | 1000G project |

* Genome assembly version=hg19.

ICGC: International Cancer Genome Consortium.

Firehose: The Broad Institute GDAC Firehose Portal.

**Table S2. Features used for constructing DGAT-cancer.**

See Additional file 2: Table S2.

**Table S3. Numbers of known cancer driver genes downloaded from Cancer Gene Census, OncoKB and IntOGen, respectively.**

| **Source** | **Cancer type** | **No. of genes** |
| --- | --- | --- |
| Cancer Gene Census | All | 723 |
| OncoKB | All | 1,064 |
| IntOGen | BLCA | 78 |
|  | BRCA | 99 |
|  | CESC | 45 |
|  | COAD | 72 |
|  | GBM | 35 |
|  | HNSC | 62 |
|  | LGG | 38 |
|  | LUAD | 42 |
|  | STAD | 35 |
|  | PCPG | 9 |
|  | TGCT | 9 |
|  | THCA | 40 |

**Table S4. Sample size of gene expression data for 12 types of cancer tissue.**

| **Cohort** | **Tumor** | **Paracancer** | **Total** |
| --- | --- | --- | --- |
| BLCA | 407 | 19 | 426 |
| BRCA | 1,104 | 114 | 1,218 |
| CESC | 305 | 3 | 308 |
| COAD | 288 | 41 | 329 |
| GBM | 167 | 5 | 172 |
| HNSC | 522 | 44 | 566 |
| LGG | 530 | 0 | 530 |
| LUAD | 517 | 59 | 576 |
| STAD | 415 | 35 | 450 |
| PCPG | 187 | 0 | 187 |
| TGCT | 156 | 0 | 156 |
| THCA | 513 | 59 | 572 |
| All | 5,111 | 379 | 5490 |

**Table S5. Data sources of mutations used in topological analysis.**

| **Cohort** | **No. of samples** | **XENA: RSEM_NORMCOUNT** | **FIREHOSE MAF FILE** |
| --- | --- | --- | --- |
| BLCA | 129 | TCGA.BLCA.sampleMap/HiSeqV2(version:2017-10-13) | Level_3.2016012800.0.0 |
| BRCA | 981 | TCGA.BRCA.sampleMap/HiSeqV2(version:2017-10-13) | Level_3.2016012800.0.0 |
| CESC | 193 | TCGA.CESC.sampleMap/HiSeqV2(version:2017-10-13) | Level_3.2016012800.0.0 |
| COAD | 267 | TCGA.COAD.sampleMap/HiSeqV2(version:2017-10-14) | xena-TCGA hub-: somatic mutation- MC3 public version |
| GBM | 149 | TCGA.GBM.sampleMap/HiSeqV2(version:2017-10-13) | Level_3.2016012800.0.0 |
| HNSC | 279 | TCGA.HNSC.sampleMap/HiSeqV2(version:2017-10-13) | Level_3.2016012800.0.0 |
| LGG | 527 | TCGA.LGG.sampleMap/HiSeqV2(version:2017-10-13) | Level_3.2016012800.0.0 |
| LUAD | 230 | TCGA.LUAD.sampleMap/HiSeqV2(version:2017-10-13) | Level_3.2016012800.0.0 |
| PCPG | 184 | TCGA.PCPG.sampleMap/HiSeqV2(version:2017-10-13) | Level_3.2016012800.0.0 |
| STAD | 272 | TCGA.STAD.sampleMap/HiSeqV2(version:2017-10-13) | Level_3.2016012800.0.0 |
| TGCT | 155 | TCGA.TGCT.sampleMap/HiSeqV2(version:2017-10-13) | Level_3.2016012800.0.0 |
| THCA | 403 | TCGA.THCA.sampleMap/HiSeqV2(version:2017-10-13) | Level_3.2016012800.0.0 |

| **Cohort** | **freq_threshold** | **subsampling** | **num_complex** | **var_threshold** | **min_interval, max_interval, interval_step** | **min_percent_overlap, max_percent_overlap, percent_step** |
| --- | --- | --- | --- | --- | --- | --- |
| BLCA | 0.001 | NA | 42 | 10000 | 20,80,10 | 60,85,5 |
| BRCA | 0.001 | NA | 42 | 10000 | 20,80,10 | 60,85,5 |
| CESC | 0.001 | NA | 42 | 10000 | 20,80,10 | 60,85,5 |
| COAD | 0.001 | NA | 42 | 10000 | 20,80,10 | 60,85,5 |
| GBM | 0.001 | NA | 35 | 10000 | 20,80,10 | 65,85,5 |
| HNSC | 0.001 | NA | 42 | 10000 | 20,80,10 | 60,85,5 |
| LGG | 0.001 | NA | 42 | 10000 | 20,80,10 | 60,85,5 |
| LUAD | 0.001 | NA | 42 | 10000 | 20,80,10 | 60,85,5 |
| PCPG | 0.001 | NA | 42 | 10000 | 20,80,10 | 60,85,5 |
| STAD | 0.001 | NA | 42 | 10000 | 20,80,10 | 60,85,5 |
| TGCT | 0.001 | NA | 42 | 10000 | 20,80,10 | 60,85,5 |
| THCA | 0.001 | NA | 42 | 10000 | 20,80,10 | 60,85,5 |

**Table S6. Arguments used in TDAmapper.**

**Table S7. Number of genes characterized by different kinds of feature.**

See Additional file 3: Table S7.

**Table S8. Number of genes involved in cancer driver prediction performed by four methods.**

| **Cohort** | **DGAT-cancer** | **MutSigCV** | **OncodriveFML** | **OncodriveCLUSTL** | **Share total^a^** | **Combine total^b^** |
| --- | --- | --- | --- | --- | --- | --- |
| BLCA | 4,164 | 18,862 | 17,744 | 20,098 | 3,842 | 21,478 |
| BRCA | 2,797 | 18,862 | 17,368 | 20,098 | 2,565 | 21,421 |
| CESC | 1,518 | 18,862 | 16,665 | 20,098 | 1,359 | 21,393 |
| COAD | 6,750 | 18,862 | 18,737 | 20,098 | 6,169 | 21,589 |
| GBM | 959 | 18,862 | 15,633 | 20,098 | 899 | 21,350 |
| HNSC | 2,685 | 18,862 | 16,963 | 20,098 | 2,486 | 21,412 |
| LGG | 223 | 18,862 | 13,096 | 20,098 | 208 | 21,324 |
| LUAD | 5,164 | 18,862 | 17,853 | 20,098 | 4,763 | 21,500 |
| STAD | 5,784 | 18,862 | 18,159 | 20,098 | 5,330 | 21,532 |
| PCPG | 2,164 | 18,862 | 2,492 | 20,098 | 1,793 | 21,541 |
| TGCT | 5,588 | 18,862 | 6,912 | 20,098 | 5,241 | 21,546 |
| THCA | 20,530 | 18,862 | 6,542 | 20,098 | 6,157 | 23,292 |

* When predicting cancer drivers for PCPG, TGCT and THCA, DGAT-cancer only used expression-based features.

a: genes predicted by all four methods: DGAT-cancer, MutSigCV, OncodriveFML and OncodriveCLUSTL.

b: genes predicted by any one of the four methods.

**Table S9. Number of positive and negative genes in calculating AUPRC when comparing four methods.**

| **Cohort** | **Positive** | **Negative** |
| --- | --- | --- |
| BLCA | 1,171 | 6,306 |
| BRCA | 1,170 | 6,258 |
| CESC | 1,168 | 6,235 |
| COAD | 1,173 | 6,401 |
| GBM | 1,168 | 6,198 |
| HNSC | 1,172 | 6,254 |
| LGG | 1,170 | 6,179 |
| LUAD | 1,169 | 6,327 |
| STAD | 1,175 | 6,354 |

**Table S10. Number of significant genes predicted by the four methods.**

| **Method**  **Cancer** | **DGAT-cancer** | **MutSigCV** | **OncodriveFML** | **OncodriveCLUSTL** |
| --- | --- | --- | --- | --- |
| BLCA | 159 | 12 | 94 | 9 |
| BRCA | 100 | 106 | 59 | 31 |
| CESC | 133 | 8 | 35 | 7 |
| COAD | 200 | 8 | 106 | 63 |
| GBM | 103 | 14 | 7 | 6 |
| HNSC | 158 | 16 | 50 | 154 |
| LGG | 36 | 16 | 11 | 8 |
| LUAD | 57 | 15 | 48 | 25 |
| STAD | 142 | 53 | 30 | 21 |
| PCPG | 34 | 4 | 0 | 96 |
| TGCT | 80 | 8 | 5 | 344 |
| THCA | 288 | 11 | 1 | 3 |

* MutSigCV: genes with $q<0.05$; OncodriveFML: genes with Q_VALUE < 0.05; OncodriveCLUSTL: genes with Q_ANALYTICAL < 0.05. When predicting cancer drivers for PCPG, TGCT and THCA, DGAT-cancer only used expression-based features.

**Table S11. Number of positive and negative genes in calculating AUPRC when comparing six methods.**

| **Cohort** | **Positive** | **Negative** |
| --- | --- | --- |
| BLCA | 1,171 | 7988 |
| BRCA | 1,170 | 7933 |
| CESC | 1,168 | 7911 |
| COAD | 1,173 | 8079 |
| GBM | 1,168 | 7870 |
| HNSC | 1,172 | 7934 |
| LGG | 1,170 | 6179 |
| LUAD | 1,169 | 8003 |
| STAD | 1,175 | 8032 |
| PCPG | 1,168 | 6175 |
| TGCT | 1,169 | 6,181 |
| THCA | 1,176 | 7,846 |

**Table S12. Novel cancer drivers of 9 cancer types predicted by DGAT-cancer.**

See Additional file 4: Table S12.

#### Supplementary Figures

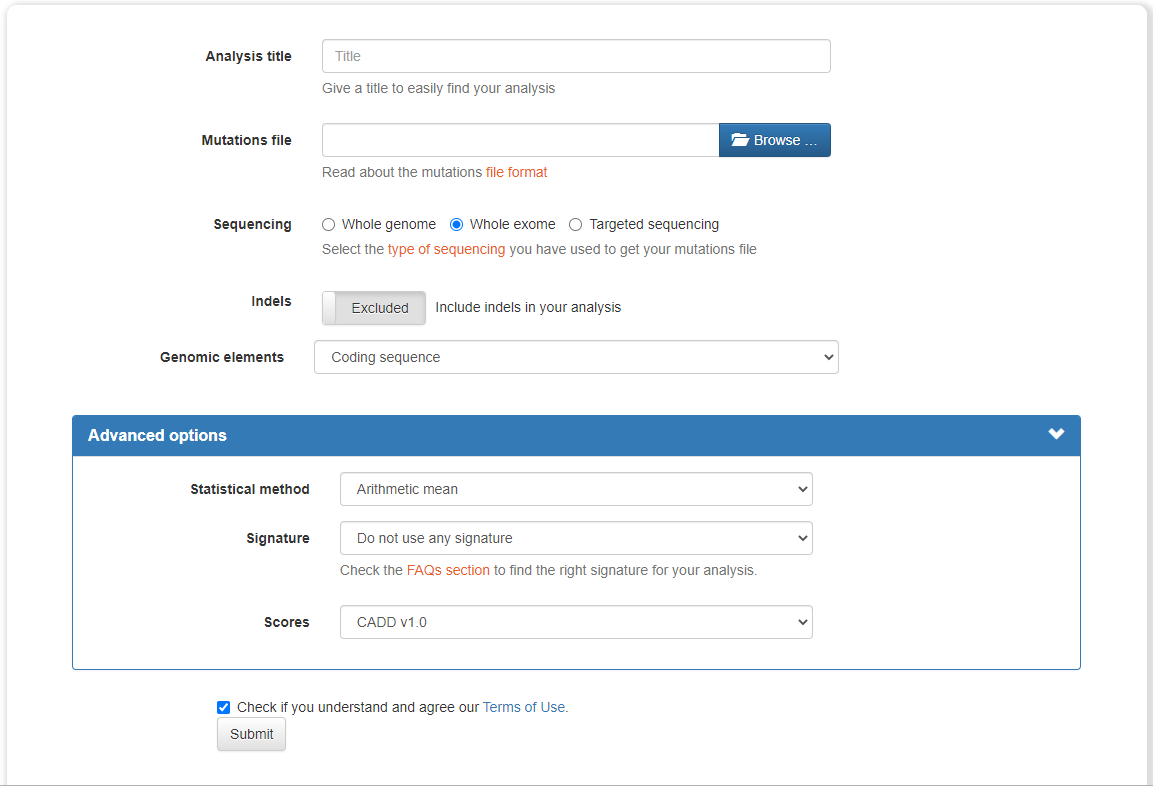

**Fig. S1. Parameters used for online OncodriveFML.**

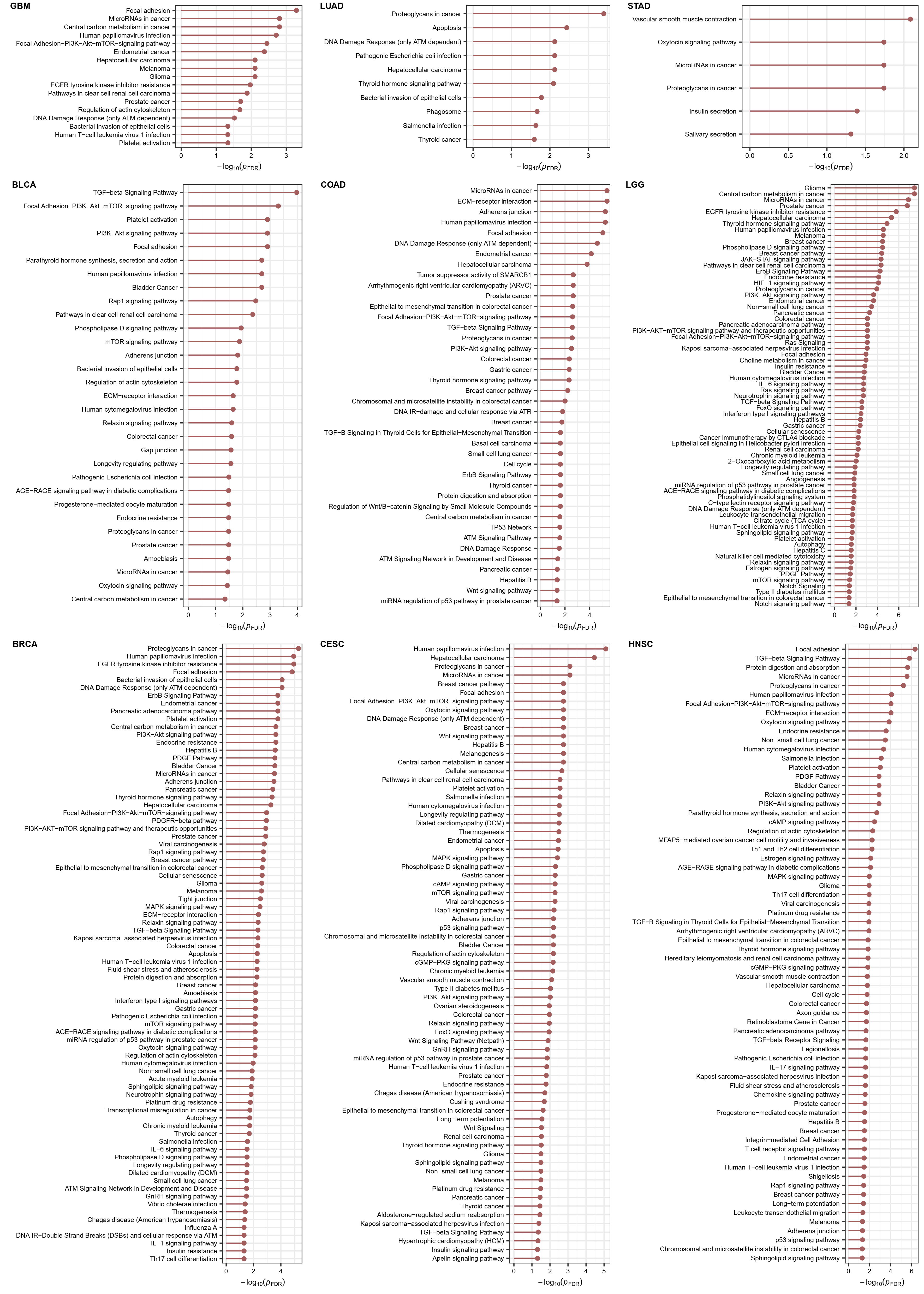

**Fig. S2. Enrichment of predicted cancer drivers in KEGG pathways.**

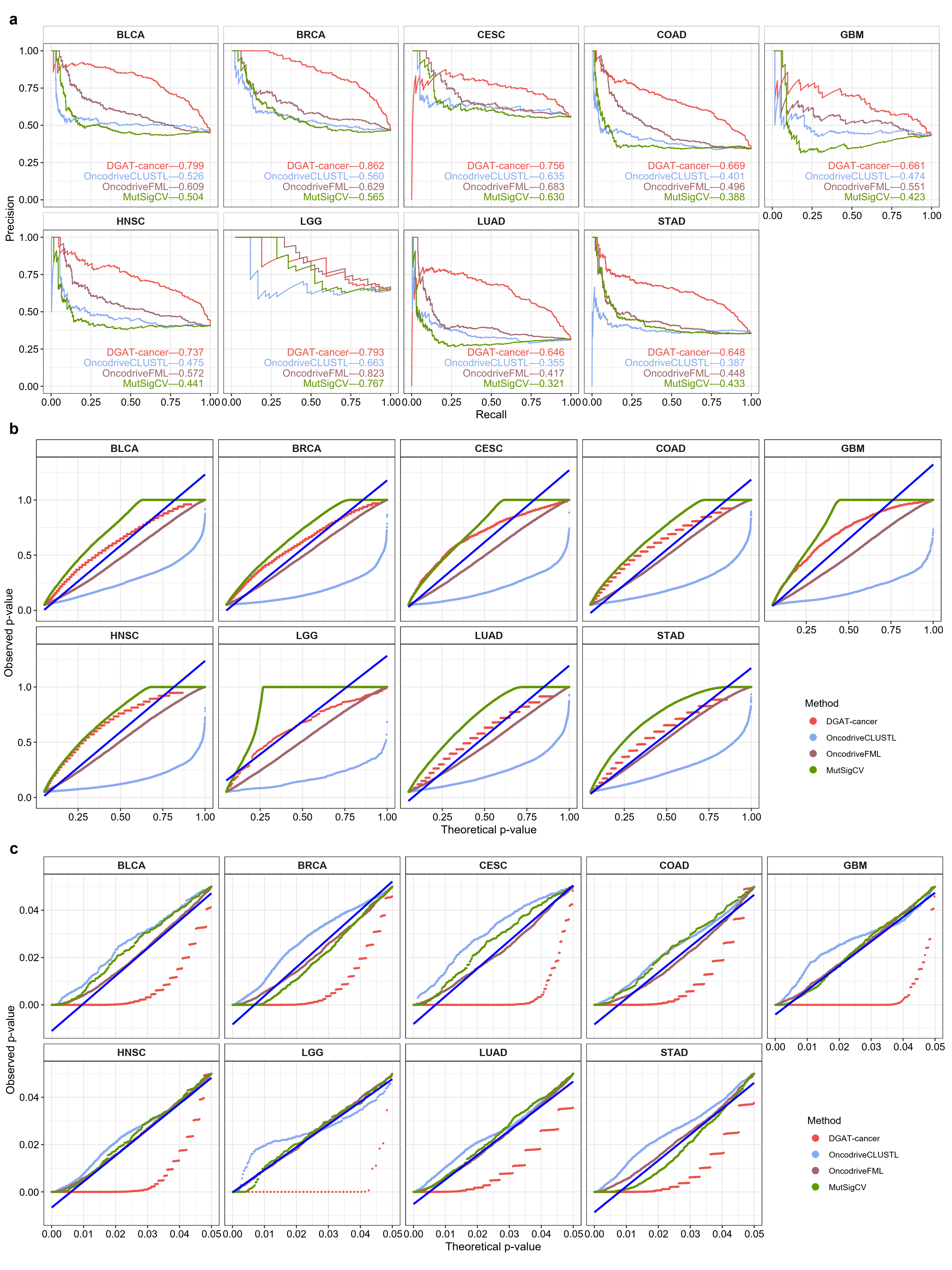

**Fig. S3.** **Comparison of DGAT-cancer with other methods with respect to their prediction of cancer drivers. a** Comparing AUPRC of the four methods for predicting cancer driver genes. The prediction was performed for genes that been predicted by all four methods. **b** Quantile–quantile plots of the P-values for genes predicted as non-cancer drivers by different methods. Observed P-values of genes predicted by different methods are compared with the expected P-values obtained from a uniform distribution (blue). **c** Quantile–quantile plots of the P-values for genes predicted as cancer drivers by the four methods.

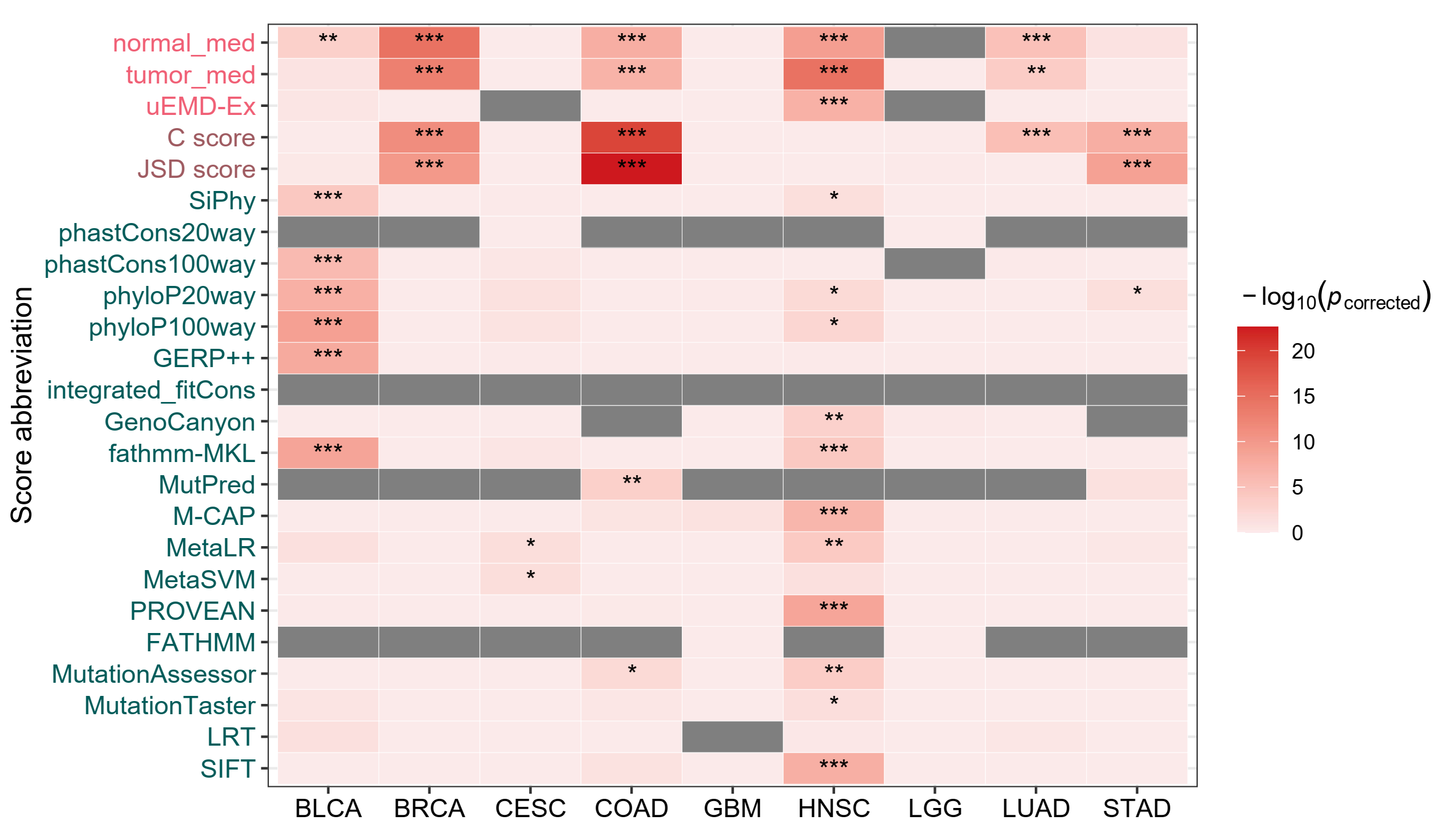

**Fig. S4. Comparison between genes missed by DGAT-cancer and genes missed by other methods on feature scores used in DGAT-cancer.** The *p*-values were obtained through Wilcoxon rank sum test and were corrected by Bonferroni. “*” means $p_{corrected}<0.05$, “**” means $p_{corrected}<0.001$, “***” means $p_{corrected}<0.0001$. Features colored in grey were filtered out in the Laplace selection and thus not used in prediction. The description of each feature is shown in Additional file 3: Table S7.

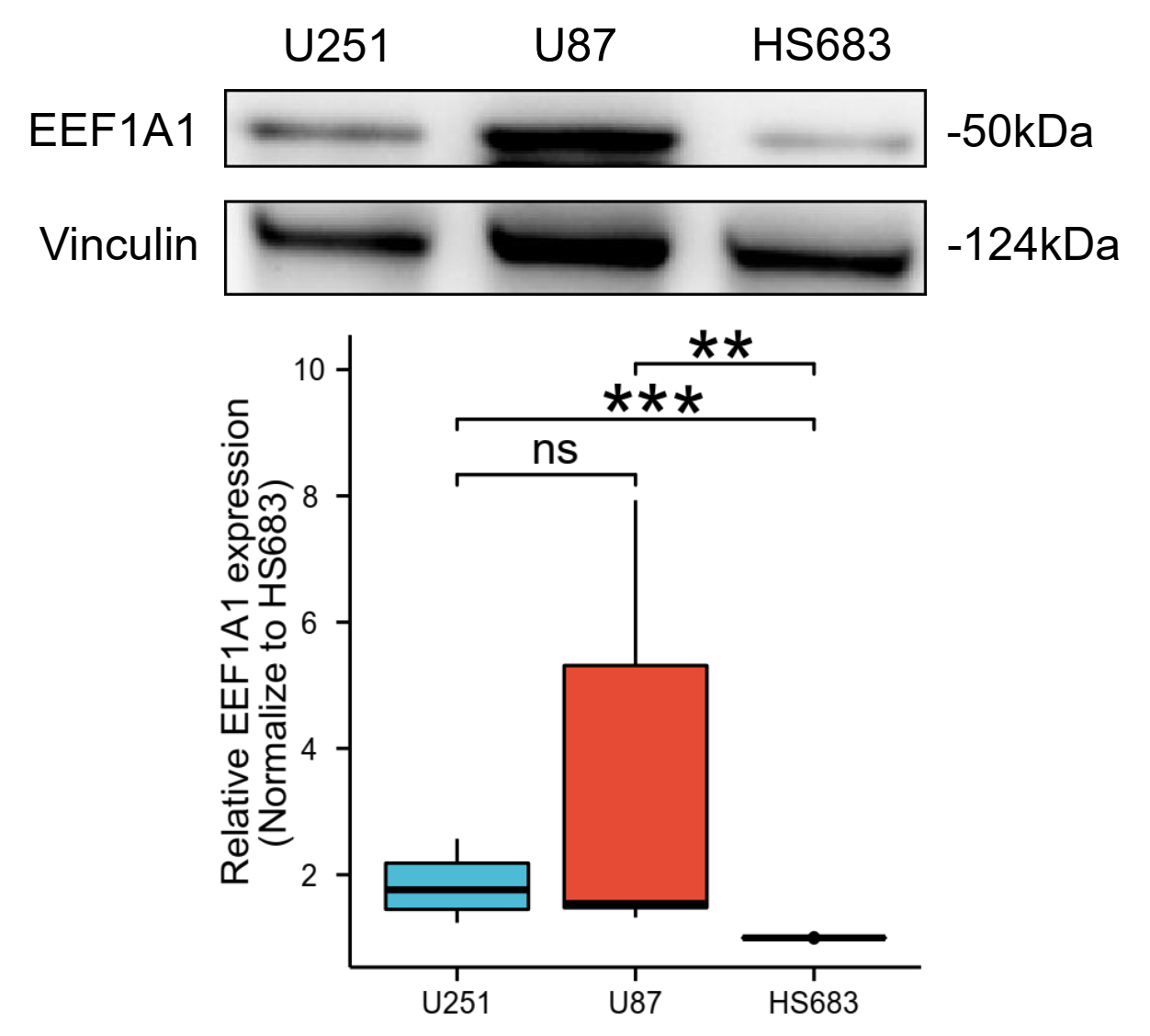

**Fig. S5.** Western blot and the quantitative expression results (n=8) of *EEF1A1* in GBM cells (U87, U251) and LGG cells (Hs683) (Kruskal-Wallis test). *p< 0.05; **p < 0.01; ***p < 0.001.

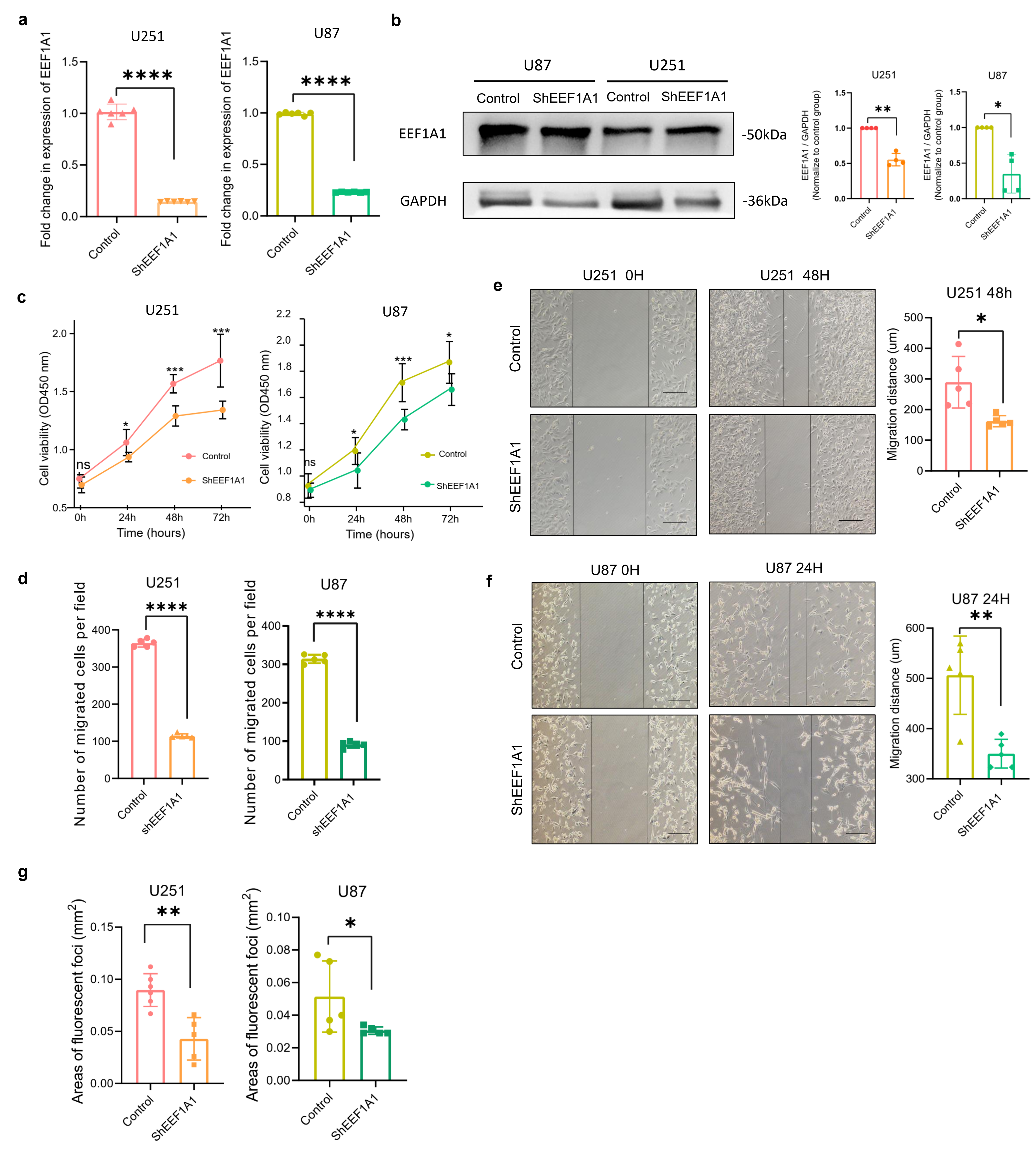

**Fig. S6. Experimental validation of roles of *EEF1A1* in GBM. a** Lentiviral knockdown of *EEF1A1* expression by shRNA in U251 and U87 cells. The efficacy of *EEF1A1* knockdown was confirmed by real-time PCR. **b** The efficacy of *EEF1A1* knockdown was confirmed by Western blot. **c** Cell viability of control and shEEF1A1 knockdown U251 and U87 cells was measured by CCK-8. N=8. **d** Quantitation of relative cell migration of (Fig.5f). **e** Scratch-wound healing assay. A significantly shorter migration distance was observed in U251 cells (*n*=5) with *EEF1A1* knockdown. **f** Scratch-wound healing assay. A significantly shorter migration distance was observed in *EEF1A1* knockdown U87 cells. N=5, Scale bar=200μm. **g** Quantitation of the areas of fluorescent foci in (Fig. 5g). * p <0.05, ** p <0.01, *** p <0.001, **** p <0.0001.

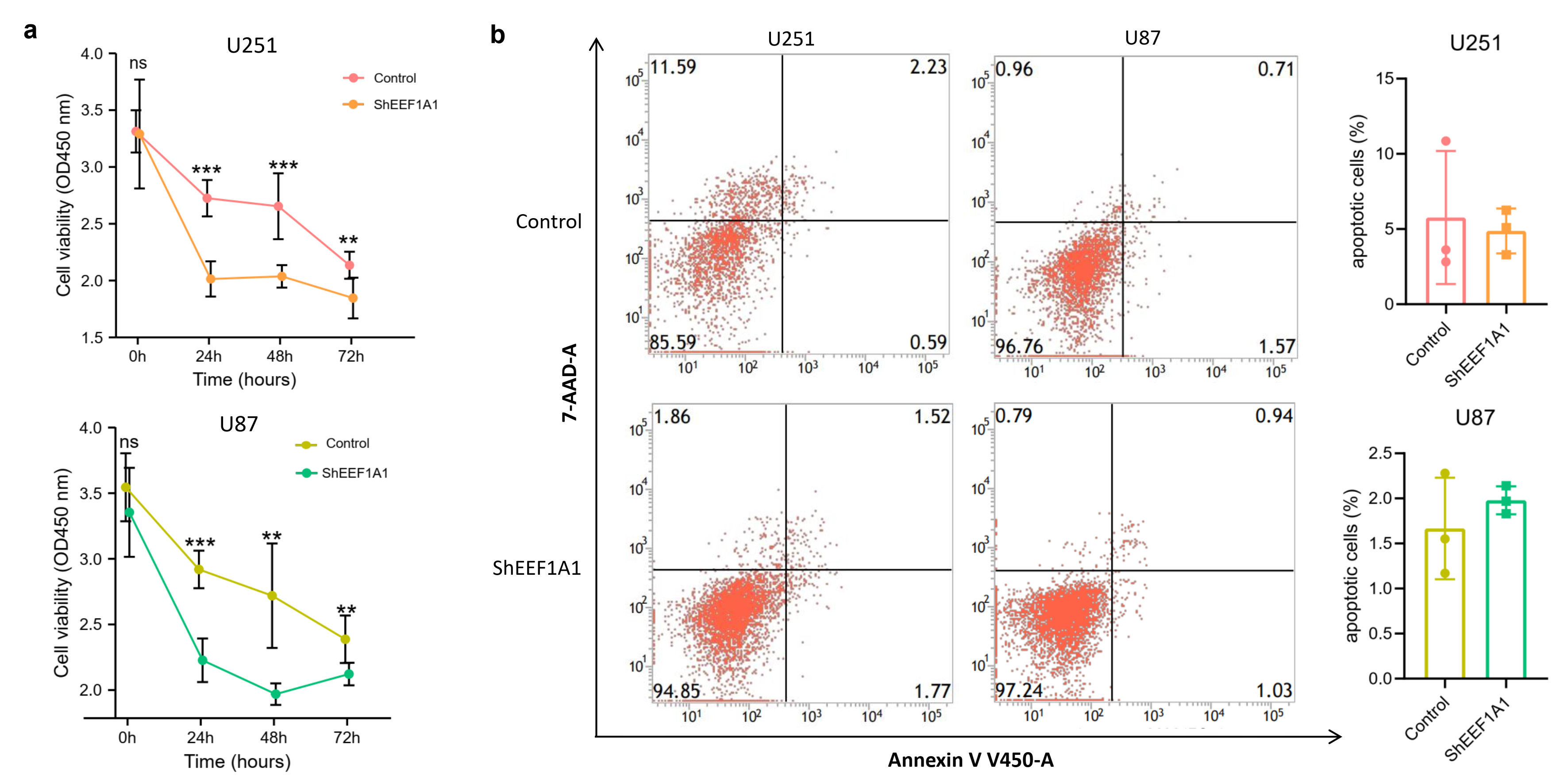

**Fig. S7. a** Culturing U251 and U87 cells in each group with 200μM TMZ, we observed that the cell viability of the EEF1A1-knockdown group was significantly lower than that of the control group according to CCK-8 assays. N=6. **b** The flow cytometry provided a measure of the proportion of apoptotic cells in the EEF1A1-knockdown and control groups in U251 and U87 cells. * p <0.05, ** p <0.01, *** p <0.001, **** p <0.0001.
